# Lagged climate-driven range shifts at species’ leading, but not trailing, range edges revealed by multispecies seed addition experiment

**DOI:** 10.1101/2023.10.25.563861

**Authors:** Katie J.A. Goodwin, Nathalie I. Chardon, Kavya Pradhan, Janneke Hille Ris Lambers, Amy L. Angert

## Abstract

Climate change is causing many species’ ranges to shift upslope to higher elevations as species track their climatic requirements. However, many species have not shifted in pace with recent warming (i.e., ‘range stasis’), possibly due either to demographic lags or microclimatic buffering. The ‘lagged-response hypothesis’ posits that range stasis disguises an underlying climatic sensitivity if range shifts lag the velocity of climate change due to slow colonization or mortality. Alternatively, the ‘microclimatic buffering hypothesis’ proposes that small-scale variation within the landscape, such as canopy cover, creates patches of suitable habitat within otherwise unsuitable macroclimates that create climate refugia and buffer range contractions. To test these two hypotheses, we combined a large seed addition experiment of 25 plant species across macro- and micro-scale climate gradients with local herbaria records to compare patterns of seedling recruitment relative to adult ranges and microclimate in the North Cascades, USA. Despite high species-to-species variability in recruitment, community-level patterns supported the lagged response hypothesis, with a mismatch between where recruitment vs. adults occur. On average, the seedling recruitment optimum shifted from the adult climatic range centre to historically cooler, wetter regions and many species recruited beyond their cold (e.g., leading) range edge. Meanwhile, successful recruitment at warm and dry edges, despite recent climate change, suggests that macroclimatic effects on recruitment do not drive trailing range dynamics. By contrast, we were unable to detect evidence of microclimatic buffering due to canopy cover. Combined, our results suggest apparent range stasis in our system is a lagged response to climate change at the cool ends of species ranges, with range expansions likely to occur slowly or in a punctuated fashion.

## Introduction

Ongoing climate change should be shifting species’ distributions. At leading edges (e.g., cold, high elevations and latitudes), previously abiotically limiting conditions should become suitable, allowing individuals to disperse and colonize beyond their leading edge. Meanwhile, at trailing edges (e.g., warm, low elevations and latitudes), increased climatic stress should drive mortality and local extinctions, retracting the range. Combined, these processes should cause upward range shifts to higher elevations and latitudes. Although many species are shifting their ranges upslope (Chen et al., 2011; Rumpf et al., 2018), range shifts are highly variable, with many ranges remaining stagnant (Rapacciuolo et al., 2014; Wilson, 2017). In many cases, it remains unclear what this frequently observed range stasis indicates for species’ sensitivity to climate change.

Range stasis in a changing climate could indicate a lack of climatic sensitivity. Alternatively, range stasis can belie sensitivity to climate change where range shifts lag the pace of atmospheric warming (hereafter termed the “lagged response hypothesis”). Ongoing range shifts can proceed at slow, undetected rates, particularly for long-lived species, as methods for detecting range shifts are often based on adult occurrence (e.g., species distribution models, Elith et al., 2010; historical plot re-surveys, Wilson, 2017). Beyond leading edges, expansion into newly macroclimatically suitable areas can be limited by dispersal (Zimmer et al., 2018), microsite availability (Goodwin & Brown, 2023), resident competition (Solarik et al., 2020), seed predators (Crofts & Brown, 2020) or a lack of mutualist partners (Benning & Moeller, 2021). These bottlenecks can create ‘colonization credits’ (Jackson & Sax, 2010) where colonization into newly suitable habitat beyond leading edges is likely but has yet to be realized. Meanwhile, at trailing edges, long-lived and more climate-tolerant adults can persist in newly climatically unsuitable regions where more climate-sensitive seedlings can no longer recruit (Davis & Gedalof, 2018; Kueppers et al., 2017). This delayed mortality creates extinction debts (Jackson & Sax, 2010) and future range contractions (Dullinger et al., 2012). In addition to shifts at edges, the geographic location of species’ climatic optima can shift to redistribute abundances upslope. However, shifts in optima can remain undetected due to the same processes described above, creating a disequilibrium between the central tendencies of the present locations of adults and seedlings (Lenoir et al., 2009). In short, the lagged response hypothesis posits that lags between demography and climate change cause a disequilibrium between where species exist and their suitable climates, creating misleading range stasis (Fig. 1A).

**Figure 1.**
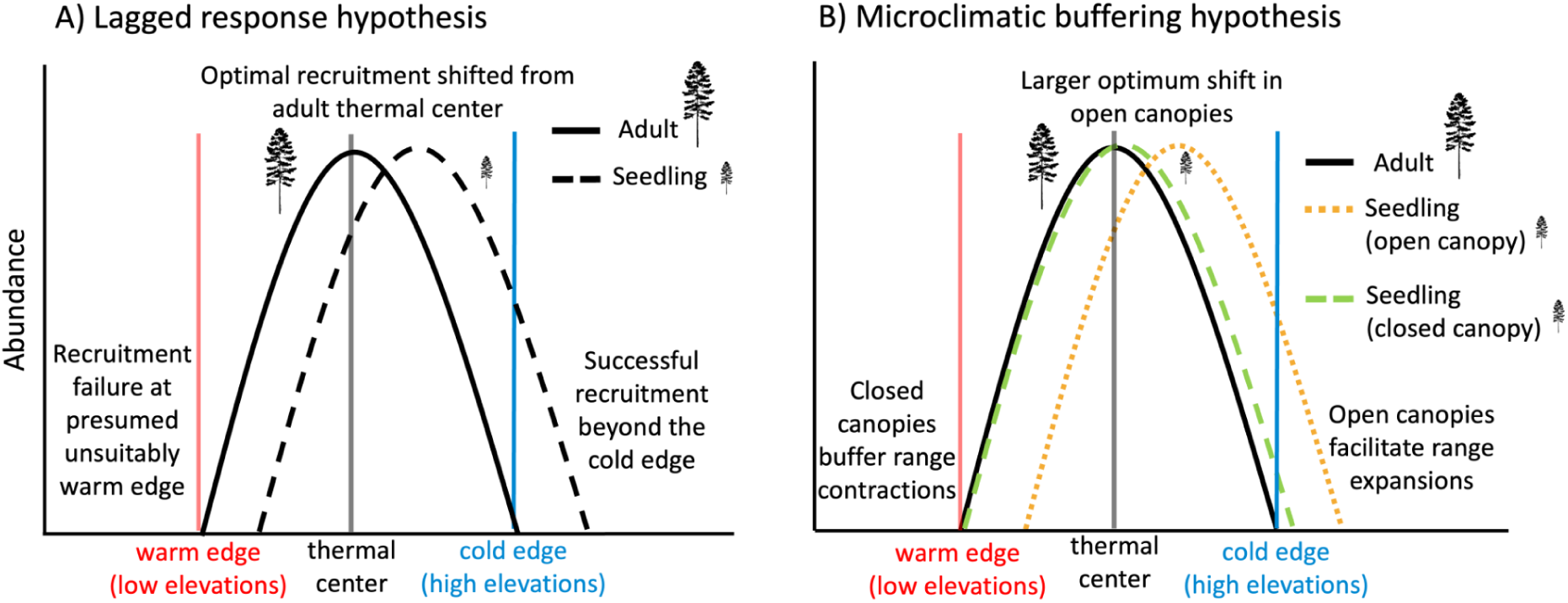
Predicted seedling recruitment (from seed addition experiment) and adult occurrence patterns across thermal ranges (from herbaria records 1980-2022) for the (A) lagged response and (B) microclimatic buffering hypotheses.

The lagged response hypothesis is not the only explanation for apparent range stasis. Microclimate can be highly variable within a heterogenous landscape (Ford et al., 2013), creating pockets of suitable microsites. The presence of suitable microclimates can allow for persistence in newly unsuitable macroclimates at the trailing edge, buffering range contractions (hereafter termed the microclimatic buffering hypothesis; De Frenne et al., 2013; Maclean & Early, 2023). For example, increased shade in denser tree canopies creates cooler and less variable ground temperatures for understory plants compared to the broader landscape (Frey et al., 2016; Vinod et al., 2023). These cooler microhabitats can protect species from atmospheric warming, allowing individuals to persist in unsuitably warm macroclimates within the range (Maclean & Early, 2023; Sanczuk et al., 2023). Meanwhile, exposed, open canopy gaps that experience more intense macroclimatic change (Sanczuk et al., 2023) may create hot spots for establishment at leading edges and facilitate upward range expansion (Tourville et al., 2022; Fig 1B).

The lagged response and microclimatic buffering hypotheses have different implications for species’ vulnerability to climate change. The lagged response hypothesis means species are sensitive to climate change, especially as climate change accelerates, while the microclimatic buffering hypothesis implies species might be somewhat resistant to climate change if they can persist in microrefugia. Both processes contribute to discrepancies between predicted and observed range shifts (Dullinger et al., 2012; Maclean & Early, 2023), but their relative importance is uncertain. Lagged responses and microclimatic buffering are revealed by a mismatch between where new individuals now recruit (i.e., seedling range) and where adults, which established under historically cooler climates, remain extant (i.e., adult range; Fig. 1). Lagged responses are the likely driver of this mismatch when discrepancies between the adult and seedling range occur in all microhabitats across the range. By contrast, microclimatic buffering is the more likely driver when seedling and adult mismatches only occur in exposed, but not buffered, microclimates. To test both hypotheses, we executed a seed addition experiment of 25 native plant species (6 trees, 9 shrubs, 8 forbs, and 2 graminoids; Table S1) across macro- and micro-scale climatic gradients in the North Cascades mountains, WA, USA to quantify how seedling recruitment (i.e., the process of germination and early establishment) varies with microclimate and across adult ranges (defined from herbaria records).

If findings support the lagged response hypothesis for thermal ranges, we predict: (1) optimal seedling recruitment has shifted from the thermal centre of adults to cooler regions, indicating an uphill shift in the location of thermal optima; (2) successful recruitment beyond leading, cold edges of adults, indicating that if dispersal limitations are overcome, species can recruit beyond their cold edge; and (3) recruitment failure at trailing, warm edges, indicating newly unsuitable conditions for recruitment within the adult range (Fig. 1A). Leading and trailing edges can also be driven by precipitation (Cahill et al., 2014; Crimmins et al., 2011). We test for potential lags in precipitation-driven range shifts via the same mechanisms described above, though our study system is experiencing complex precipitation changes involving changes in precipitation type and seasonality rather than with elevation (Mote & Salathé, 2010; PCIC, 2021). If findings support the microclimatic buffering hypothesis, we predict (4) seedlings in open canopies will exhibit lagged recruitment patterns, similar to the previous hypothesis, while recruitment in closed canopies will align with adult range position because closed canopies buffer range contractions while open canopies facilitate range expansions (Fig. 1B). Regions of high initial recruitment do not necessarily translate to high establishment if later life stages exhibit different climate sensitivities (Donohue et al., 2010). Therefore, we monitored seedling growth and survival for five years to test if later demographic processes alter initial recruitment patterns. Combined, findings strengthen our understanding of how species distributions will respond to climate change by testing the relative importance of demographic lags and microclimatic buffering (Dullinger et al., 2012; Maclean & Early, 2023), two mechanisms that have very different conservation implications.

## Materials and methods

### Seed addition experiment across macro- and micro-scale gradients

To quantify seedling ranges and test both the lagged response and microclimatic buffering hypotheses, we implemented a seed addition experiment in the North Cascades mountains, USA (study design diagram: Fig. S1). At the macroscale, we established one transect each on the wetter west side (Mount Baker National Forest) and drier east side (Okanogan National Forest) of a rain shadow gradient on the traditional lands of the Nlaka’pamux, Nooksack, Okanagan, and Methow peoples. Along each transect, we selected sites every ∼100m of elevation at 15 elevations per transect spanning 529 m to 1517 m and 794 m to 2028 m of elevation at Mount Baker and Okanogan, respectively. To test whether canopy cover buffers range shifts, we selected two sites within each elevation: one with a relatively more open canopy and one with a more closed canopy. We measured percent canopy cover with a densiometer two times a year throughout the experiment at each plot. In total, our seed addition sites spanned a gradient of temperature (1.7°C - 8.9°C mean annual temperature [MAT]), precipitation (529 - 2981 mm mean annual precipitation [MAP]), and canopy cover (4.6% - 96.5% canopy cover). Each site had three replicate plots ∼2 m from one another, each containing three 50 cm x 50 cm quadrats ∼20 cm from one another. Quadrat locations had no saplings >10 cm in diameter, no large rocks that covered more than 10% of the quadrat, and ≥ 5% cover from seedlings or understory species to ensure substrates were potentially suitable for recruitment. The three quadrats consisted of a control quadrat (no seeds added) and two quadrats with seeds added. We split species within the same family into separate quadrats to aid in species identification at the seedling stage. In total, there were 2 transects x 15 elevations x 2 canopy covers x 3 replicate plots x 3 quadrats for 540 quadrats (or 180 plots).

We selected 25 native species encompassing a variety of functional groups (Table S1). Species included congeneric and confamilial pairs with different distributional characteristics (i.e., warm/cold affiliated or wet/dry affiliated) and high regional prevalence to capture a variety of portions of species’ climatic ranges. We collected as many species’ seeds locally as possible in 2016 and purchased the rest from local nurseries. Seed source did not affect recruitment likelihood in our experiment (Fig. S3). We combined seeds into multi-species mixtures by mass, with approximately 0.25 g of seed per species per quadrat. Thus, species with small seeds were added in greater number than large-seeded species, as per capita emergence is inversely proportional to mass (Bond et al., 1999). Seeds were mixed with ∼80 mL of sand and scattered on quadrats after raking the surface lightly at the end of the 2017 growing season (September to October). To assess recruitment, we marked new seedlings with unique coloured toothpicks for each species, counted, and monitored for survival to the end of each growing season (August to September) from 2018-2022 (excluding 2021 due to US-Canada international border restrictions). We monitored only survival, not new germination, in 2022 as we were unable to reliably estimate new recruits since sites were not visited in 2021; we also presume that new recruitment was minimal five years after seed addition. We recorded seedling height for up to three randomly selected seedlings for each species in each germination cohort to capture differences in early performance following initial recruitment.

### Quantifying adult climatic ranges

To quantify adult ranges, we estimated climatic ranges from herbaria records for each species, including synonyms, from January 1, 1980 – January 31, 2022 from the Global Biodiversity Information Facility (http://data.gbif.org/; downloaded on Jan. 31, 2022). We assume herbaria records primarily represent adult life stages. We restricted our search to occurrences within the North American temperate conifer forest biome (defined by Ecoregions, 2017). The rationale was to use records from broadly similar habitats to our study system while encapsulating a sufficient sample size; filtering to records only from the North Cascades resulted in too few occurrences. We removed occurrences with high uncertainty (less than three decimal places for latitude and longitude). To calculate the climatic range of each species, we downloaded the 1981-2010 normal mean annual temperature (MAT), mean warmest month temperature (MWMT), mean annual precipitation (MAP), and May to September precipitation (MSP) for each occurrence record from gridded climate normal data (800 m x 800m) (Wang et al., 2016). This timeframe captures the mean climate that extant (or recently extant) adults of long-lived species occupy to test for climatic mismatches between adults and seedlings. While some adults from earlier herbaria records might have died by the time of the experiment, with potential local extinctions at trailing edges, our recent study by Wilson et al. (2017) found most species in our system have not shifted their adult ranges from 1983-2015. We found high correlation amongst temperature variables and amongst precipitation variables across herbaria records (Pearson’s correlation coefficient, *r* > 0.59) and study sites (*r* > 0.85). We selected MAT and MAP to describe species’ thermal and moisture ranges, respectively. MAT and MAP were uncorrelated (*r* = 0.17; Fig. S2).

To facilitate generalization across species, which differ in their climatic ranges, we created standardized thermal and precipitation range position variables to determine where each seed addition site occurred in each species’ adult climatic range. We defined the lower and upper thermal and precipitation ranges as the 2.5% and 97.5% quantiles, respectively, of the distribution of MAT and MAP herbaria occurrences for each species (climatic ranges in Table S1). We selected these quantiles to remove potentially inaccurate outliers from herbaria records. We then calculated where each site fell within each species’ adult climatic range, which was standardized for each species. We used species’ climatic midpoint to represent the thermal centre to avoid sampling bias (i.e., some climates might be oversampled in herbaria records). This assumes that the thermal optima for species are in the center of their thermal range, which could add noise to our results as some species true optima may lie in cooler or warmer regions. We calculate climatic range position using the equation:

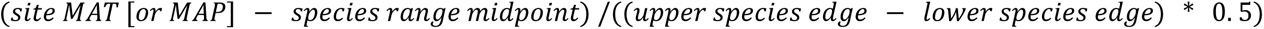

This equation created two standardized variables: thermal and precipitation adult range position, where 0 indicates the range centre, -1 is the cold or dry range edge, +1 is the warm or wet range edge, -1 to 0 is the cooler or drier half of the species’ range, and 0 to +1 are the warmer or wetter half of the species’ range. Values > 1 and < -1 indicate that sites were respectively above and below a species’ adult range (see Fig. 2 for thermal range position of sites; Fig. S4 for precipitation range position of sites).

**Figure 2.**
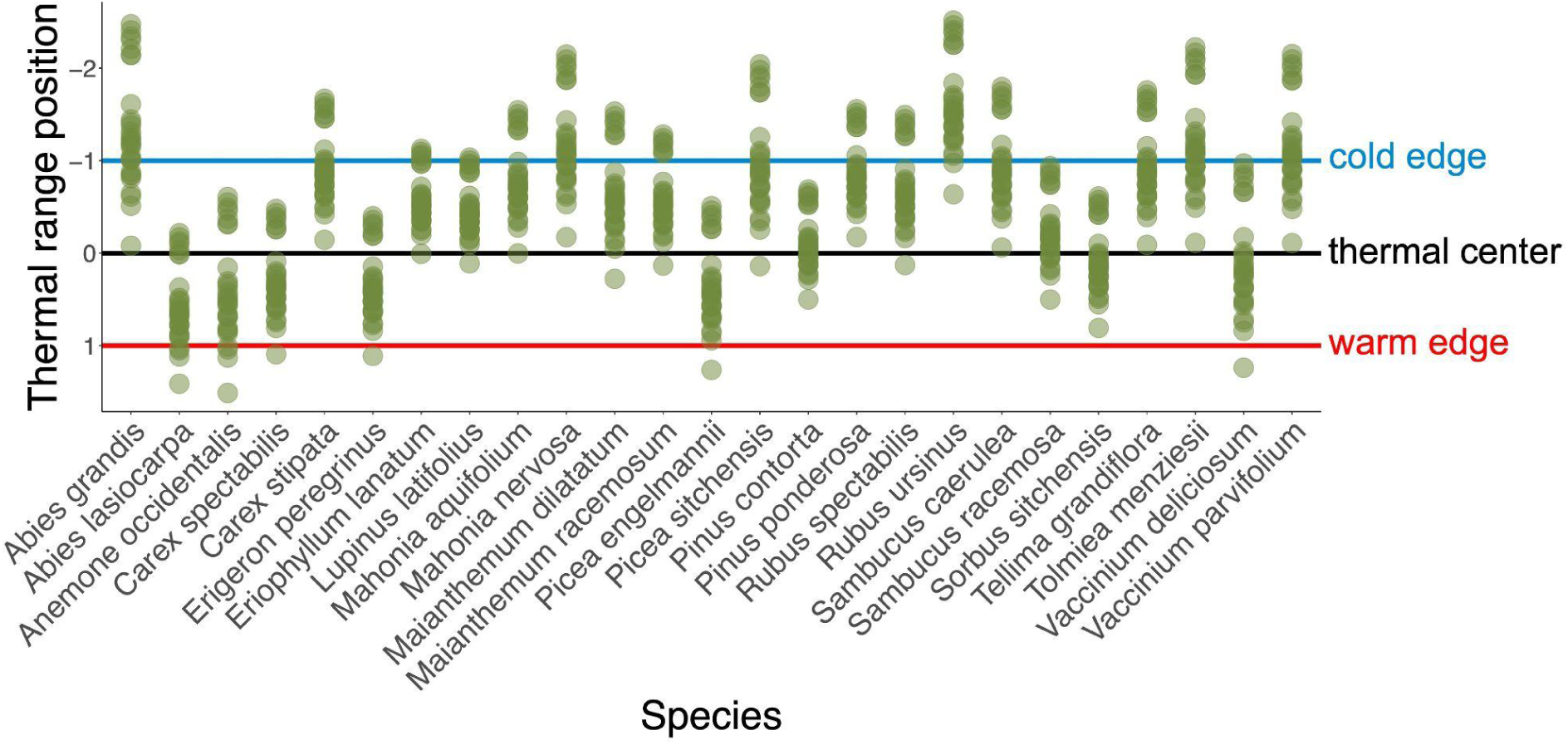
Thermal range position of each study site for each focal species. Green points indicate a thermal range position of a study site for a given species. The cold edge, thermal centre, and warm edge are respectively shown in blue, black, and red lines. Thermal range position was quantified using mean annual temperature data from 1981-2010 (Wang et al., 2016) of herbaria occurrences for each species (http://data.gbif.org/; downloaded on Jan. 31, 2022).

### Data analysis

To test both the lagged response and microclimatic buffering hypotheses, our response variable was the number of seedlings of a focal species that recruited in a given plot any year throughout the experiment (‘seedling count’). We first adjusted for any background recruitment by subtracting recruitment in a control quadrat from recruitment of that focal species in the paired seed addition quadrat. To assess whether any models were driven by the few species that had frequent germination, we also considered models with a response variable of relative recruitment normalized to the maximum recruits observed for each species in a given plot in the experiment. However, these relative recruitment models did not change any results and had poor diagnostics, so we used seedling count as the response variable for our final models. We assessed model fit and checked model assumptions for all models using scaled simulated residuals conditional on the inferred random effects using the DHARMa package (Hartig & Lohse, 2022) in the R environment version 4.2.1 (R Core Team, 2022) and followed Zuur et al’s (2009) mixed effects modelling procedure.

### Data analysis: Lagged response hypothesis

We conducted three analyses to test for macroclimatic mismatches between adult and seedling ranges (i.e., the lagged response hypothesis) at the centre, cold edge, and warm edge (Predictions 1-3; Fig. 1A). To test whether optimal seedling recruitment had shifted from the adult climatic centre (Prediction 1), we subsetted our dataset to species that recruited in at least 8 of the 180 total plots to exclude species for which we had insufficient information on recruitment across the range. Models including species with less recruitment success had poor diagnostics. 14/25 species were included in the model: *Abies lasiocarpa, Anemone occidentalis, Erigeron peregrinus, Eriophyllum lanatum, Lupinus latifolius, Mahonia aquifolium, Mahonia nervosa, Picea engelmanni, Rubus ursinus, Sorbus sitchensis, Tellima grandiflora, Tolmiea menziesii, Vaccinium deliciosum,* and *Vaccinium parvifolium*. We used a zero-inflated, negative-binomial, mixed effects model using the glmmTMB package (Brooks et al., 2017). Zero-inflated models separately consider different variables for the zero-inflation part of the model (i.e., probability of a non-zero, that we consider to explain the probability of any successful recruitment across the range) and the conditional part of the model after accounting for zero-inflation (i.e., if recruitment is possible, where across the range does optimal recruitment occur?). We primarily focus on the conditional part of the model to identify climatic range position for optimal recruitment but note if regions of the range are unsuitable for recruitment success. The model equation for both parts of the model was: seedling count ∼ thermal range position^2^ + precipitation range position^2^ + (1|genus) + (1|site/plot). We used genus rather than species to account for the phylogenetic similarity of the two *Vaccinium* species.

To test whether the peak of the quadratic relationship between recruitment and thermal range position (i.e., optimal recruitment) from the above model had shifted from the adult thermal centre (Prediction 1), we used a bootstrapping procedure. We randomly resampled the seedling data (preserving sample sizes and the nested structure of the data), refit the model 5000 times, and used outputs to determine the percentage of bootstraps with optimal recruitment in the cooler half of the climatic range (i.e., percentage of bootstraps supporting the lagged response hypothesis; Prediction 1; Fig. 1A). We also used bootstrap outputs to generate bias-corrected 95% confidence intervals of the predicted relationship (Diciccio & Romano, 1988). We conducted the same bootstrapping procedure to identify shifts in the precipitation optimum for seedling recruitment. To identify species-specific recruitment patterns across thermal ranges, we ran separate models for the 14 individual species that recruited in at least 8 plots (see online supplementary for analysis).

To determine whether seedlings successfully recruited beyond the adult cold edge, as per the lagged response hypothesis (Prediction 2), we binned thermal range position into two levels, one for sites beyond the adult cold edge (thermal range position < -1) and one for sites in the cooler half of the adult range (thermal range position 0 to -1). We subsetted data to species that successfully recruited in the experiment and for which at least two sites were beyond their cold edge (13 species; Table S1). We tested for differences in seedling count within vs beyond the cold edge using a zero-inflated mixed effects model with a negative binomial distribution (formula: seedling count ∼ cool thermal range (factor) + precipitation range position^2^ + (1|species) + (1|site/plot)). We consider non-zero recruitment beyond the adult cold edge as support for the lagged response hypothesis (Fig. 1A).

To test for recruitment failure at the warm edge as predicted under the lagged response hypothesis (Prediction 3), we defined a zone of a species’ adult range that should be unsuitably warm for seedlings, if the edge is trailing (i.e., the area between adult and seedling warm edges in Fig. 1A). Assuming that range shifts are symmetrical, such that a shift at a species’ warm edge is equal to the shift in its recruitment optimum, the presumed seedling warm edge was defined based on the shift in optimal recruitment between seedlings and adults. For example, if optimal seedling recruitment is at an adult thermal range position of -0.26, then the presumed seedling warm edge is at 1 – 0.26 = 0.74; or shifted 13% from the adult warm edge (as climatic ranges span -1 to +1). Based on this threshold, we binned thermal range position into two levels, one for sites that are presumed unsuitably warm (i.e., beyond the presumed seedling warm edge; > 0.74) and one for sites in the warm half of the range (i.e., thermal range position of 0 - 0.74). We subsetted data to species that successfully recruited in the experiment and for which at least two sites were beyond this presumed seedling warm edge (6 species; Table S1). Our model to test for differences in recruitment between the presumed unsuitably warm and warm half of the range was: seedling count ∼ warm thermal range (factor) + precipitation range position^2^ + (1|species) + (1|site/plot). We consider zero recruitment beyond the presumed seedling warm edge as support for the lagged response hypothesis (Fig. 1A). We also note which species recruited up to their adult warm edge, to have more confidence in our inferences regarding warm edge recruitment failure in case range shifts are not symmetrical. We also tested for similar lagged recruitment responses beyond both the wet and dry edges, following the same procedures described above.

### Data analysis: Microclimatic buffering hypothesis

We conducted three analyses to test the microclimatic buffering hypothesis at the range centre, cold edge, and warm edge (Prediction 4; Fig. 1B). To test if open canopies exhibit larger shifts in optima than closed canopies, we used a model similar to the model for lagged responses, but we added an interaction term between canopy cover and thermal range position (formula: seedling count ∼ canopy cover + thermal range position^2^ + canopy cover:thermal range position + transect + (1|species) + (1|site/plot)). Canopy cover was excluded in prior models due to convergence issues, so for this analysis we primarily focused on thermal rather than precipitation range position, as the effects of canopy cover on precipitation are more complex and do not lead to clear predictions of microclimatic buffering (e.g., Liu et al., 2014; von Arx et al., 2013). Indeed, our study system showed no trend in soil moisture with canopy cover (Fig. S9 A,B). The canopy cover variable was the average percent canopy cover of all measurements per plot. We subsetted to species that recruited in at least 5 plots (instead of 8 plots, as for the lagged response optimum model) to include enough data to test for an interaction. To test whether canopy cover influences recruitment at the cold and warm edges, we added an interaction term between thermal range position and canopy cover in both range edge models (formula: seedling count ∼ thermal range (factor) + precipitation range position^2^ + canopy cover + canopy cover:thermal range + (1|genus) + (1|site/plot)). We consider a p-value for any interaction term between canopy cover and range position that is significant at α = 0.05, where optimal recruitment is shifted to the cooler portion of the range in increasingly open canopies, open canopies increase recruitment beyond the cold edge, and closed canopies increase recruitment beyond the warm edge, to be support for the microclimatic buffering hypothesis (Prediction 4; orange versus green lines in Fig 1B).

### Data analysis: Establishment following initial recruitment

The models described above test hypotheses at the initial recruitment phase (i.e., whether and how many seedlings germinated and persisted to the end of their first growing season throughout the experiment). To determine whether any patterns are altered following initial recruitment, we modelled how seedling growth and survival varied across climatic ranges. For growth, we modelled plant height at the end of the first growing season using the Gaussian distribution. We standardized height by the species mean and standard deviation to account for species differences in height and removed one outlier (> 4 standardized units above the mean) (formula: height ∼ thermal range position + precipitation range position + (1|site/plot)). We separately modelled the probability of seedlings surviving to their second year, third year, and to the end of the experiment (i.e., up to five years depending on when seeds germinated) using the binomial family (formula: proportion survived ∼ thermal range position + precipitation range position + (1|site/plot)) weighted by the denominator of proportion survived. Year germinated was included in the year two survival model as it included multiple cohorts of recruits. We considered both quadratic and linear relationships for range position terms and selected models with the lowest Akaike information criterion (AIC) score. We adjust conclusions drawn from seedling count models if survival and growth models show different climatic preferences than the seedling count models (i.e., if survival and growth patterns indicate that conditions for longer-term establishment differ from the initial recruitment phase).

## Results

Overall, we observed low recruitment in the seed addition experiment. Of the 25 species added to 180 replicate plots (total 4500 species – plot combinations), 91.3% of species-plot combinations had zero recruitment from the focal species after five years. 4/25 species had complete recruitment failure across the experiment (*Carex spectabilis, Maianthemum racemosum, Maianthemum dilatatum,* and *Pinus contorta*). *Maianthemum racemosum* and *M. dilatatum* failed to germinate in laboratory germination trials (*unpublished data*), exhibit multi-year dormancy, and have strict germination requirements (Hough, 2008; Kawano et al., 2020), which provides a partial explanation for the recruitment failure of these species. Nonetheless, 21/25 species recruited (393 species-plot combinations), with a total of 2080 seedlings followed for up to 5 years (range 1 - 469 seedlings per species).

### Lagged response hypothesis

As we predicted for the lagged response hypothesis, optimal recruitment shifted to the cooler half of the range relative to the adult centre, but recruitment success at the warm end of the range indicates a lagged response only at the leading, not trailing, edge of the range (Prediction 1-3; Fig. 3; Fig. 4A; Fig. 5; Table 1). The probability of recruitment did not vary across the range (i.e., zero inflation portion of model; Fig. S7). However, the peak of the recruitment curve (i.e., conditional portion of the model; Prediction 1) was at a thermal range position of -0.26 (i.e., shifted 13% from the adult thermal centre towards the cold edge). 93.94% of bootstrap resamples had optimal recruitment in the cooler half of the range (Fig. 4A). Optimal recruitment also shifted to the wetter part of the range from the adult precipitation centre (Fig. 4B; Fig. S5; Table 1). The peak of the recruitment curve was at a precipitation range position of 0.12 (i.e., shifted 6% towards the wet edge). 94.46% of bootstrap resamples had optimal recruitment within the wetter half of the range (Fig. 4B). These general patterns only emerged when examining patterns across all species. Species-specific models had idiosyncratic responses and wide confidence intervals due to small sample sizes (Fig. S8; Table S3).

**Figure 3.**
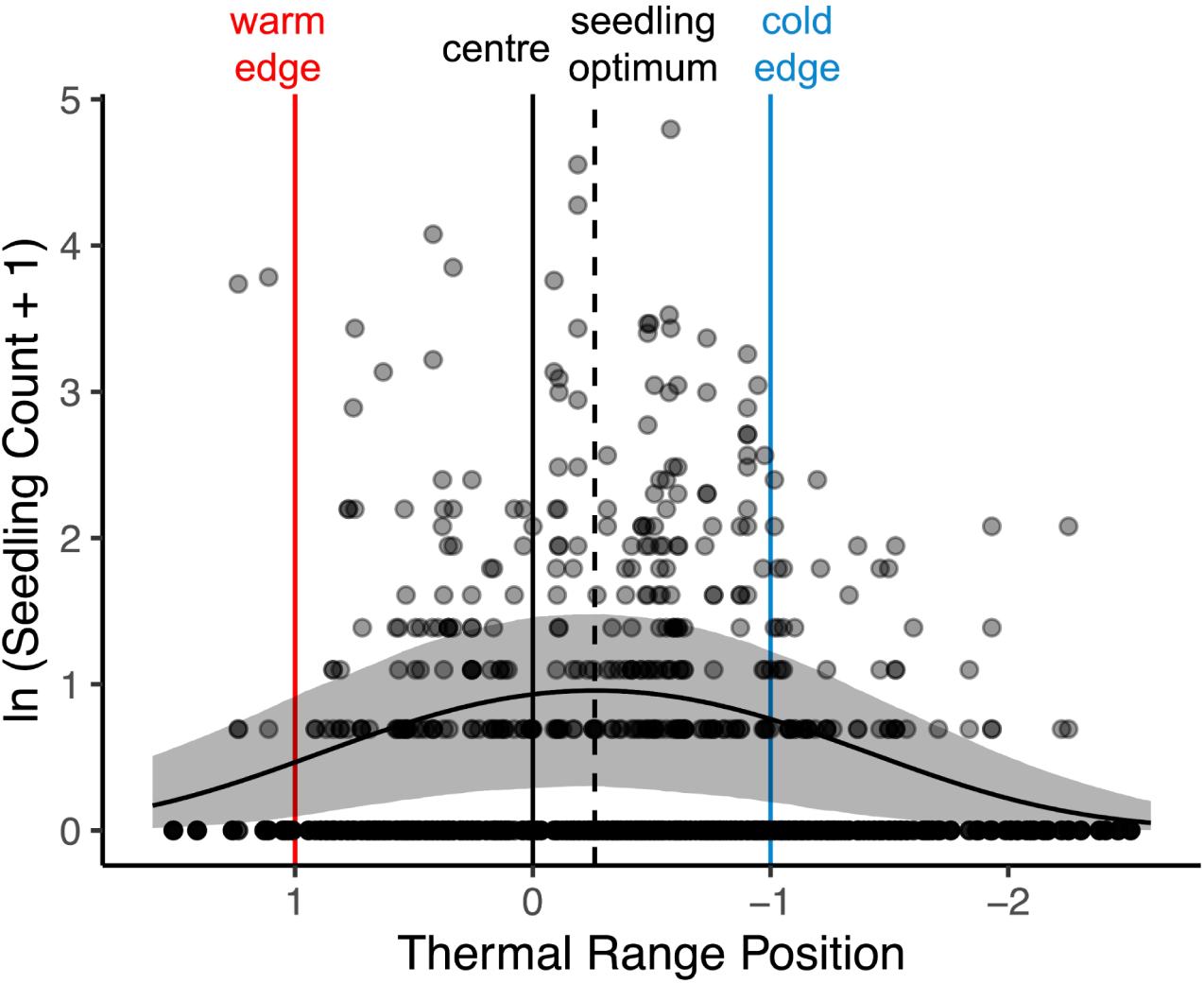
Optimal seedling recruitment (vertical dashed black line) has shifted from the adult thermal centre (vertical solid black line) to cooler regions of the range. Model-predicted recruitment as a function of thermal range position at the community level for 25 species in the North Cascades, WA. The black curve is the fitted line from the conditional portion of the zero-inflated linear model (i.e., if recruitment is possible, the peak of the curve tells us where recruitment is optimal across the range). Shading shows the bias-corrected 95% confidence interval around the fitted line derived from a bootstrapping procedure. Points represent the number of recruits for a given species in a plot. The warm edge and cold edge are respectively shown with vertical red and blue lines. Model results summarized in Table 1, and the zero component is presented in Fig. S7.

**Figure 4.**
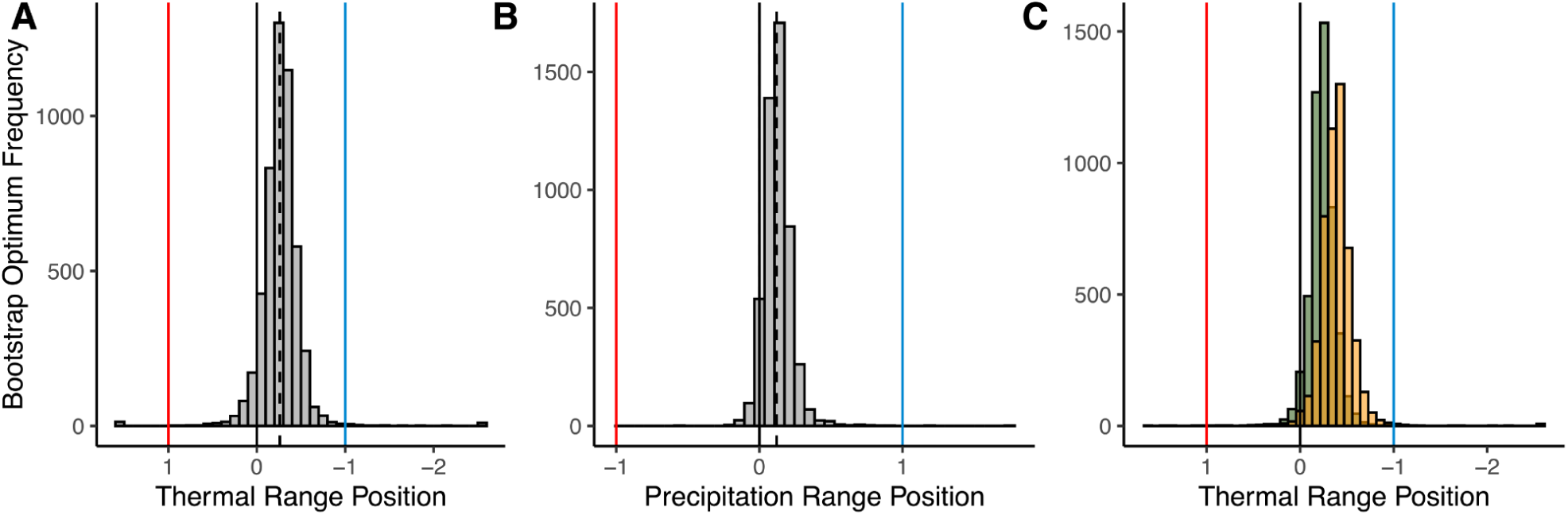
Optimal recruitment is in (A) cooler and (B) wetter regions of the range, regardless of (C) canopy cover. The distribution of optimal recruitment (i.e., peak of quadratic curve) from a bootstrapping procedure for (A) thermal range position, (B) precipitation range position, and (C) thermal range position with relatively more open and closed canopy covers. The warm/dry edges, centre, and cold/wet edges are respectively shown with vertical red, black, and blue lines. The fitted recruitment optimum is shown with a vertical dashed line. Green and orange for (C) respectively represent mean canopy cover of closed sites (88% canopy cover) and mean cover of open sites (64% canopy cover). 93.94% and 94.46% of bootstrap resamples had optimum recruitment in the cooler and wetter portion of the adult range, respectively. There was no significant difference in optimal recruitment with varying canopy cover. For full predicted recruitment relationships see Fig. 3 for thermal range, Fig. S5 for precipitation range, and Fig. S10 for canopy cover.

**Figure 5.**
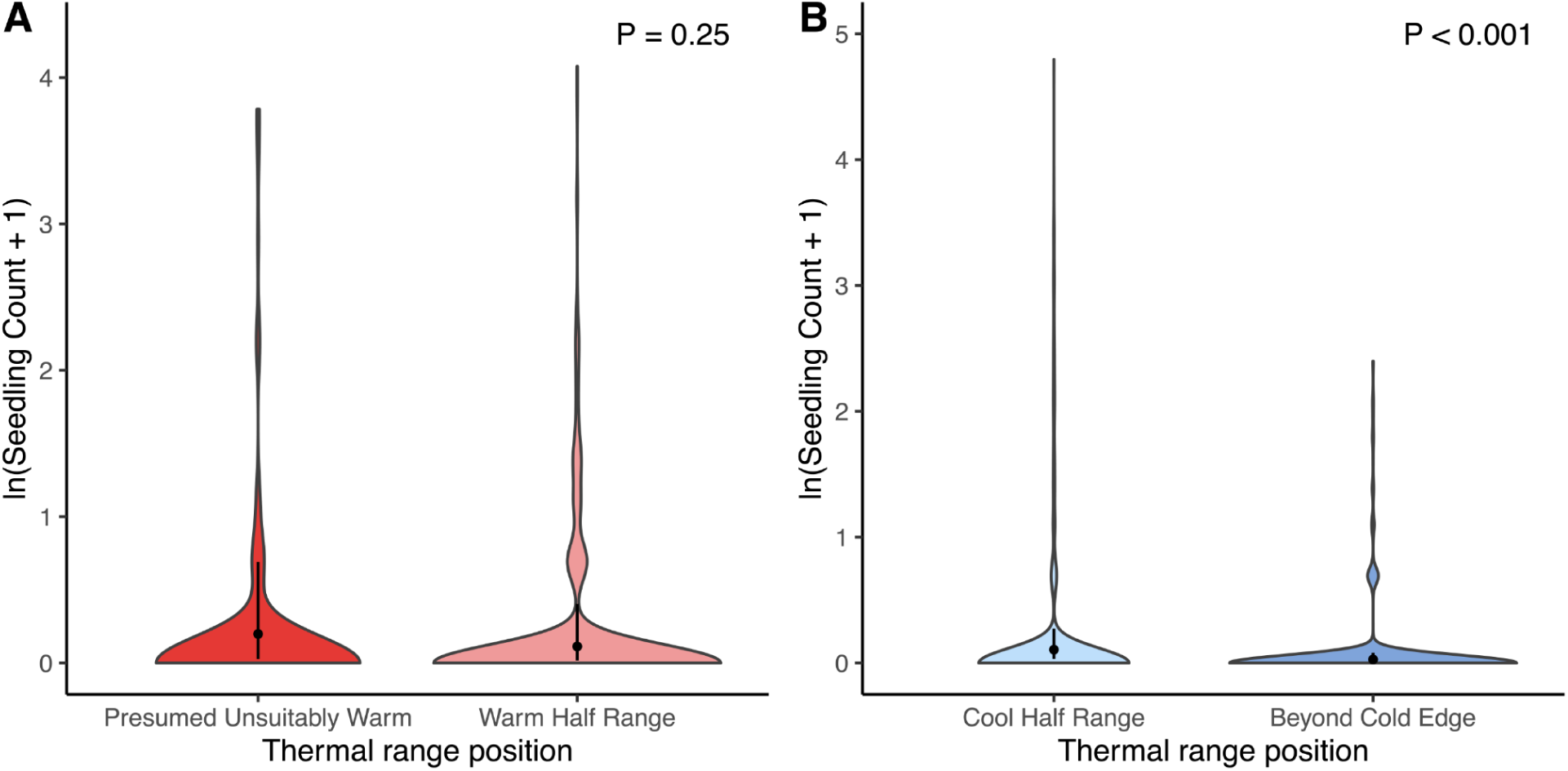
Predicted differences in recruitment between (A) presumed unsuitably warm regions of the gradient and the warm half of the range; and (B) the cool half of the range and beyond the adult cold edge. Species successfully recruited beyond both range edges. The presumed unsuitably warm bin is defined as greater than the presumed seedling warm edge (thermal range position > 0.76, based on the optimal recruitment shift between adults and seedlings). Warm half of range is a thermal range position 0 to 0.76. Beyond cold edge is a thermal range position < -1. Cool half of range is a thermal range position -1 to 0. Shaded density plot shows the distribution of number of recruits for each species in a given plot. Colour indicates thermal range bin below the presumed warm edge (dark red), within the warmer (light red) or cooler (light blue) half of the range, and beyond the cold edge (dark blue). Black points show predicted recruitment count, lines show bias-corrected 95% confidence intervals for predicted recruitment count derived from a bootstrapping procedure. Model results summarized in Table 1.

**Table 1.** Results of zero-inflated generalized mixed linear models testing whether seedling recruitment varies with climatic range as per the lagged response hypothesis for 25 species in the North Cascades, WA. The optimum model fit seedling number as a function of continuous climatic variables (“Temp.” = mean annual temperature; “Ppt.” = mean annual precipitation) describing the position of each site relative to a species’ climatic range. Edge models compared seedling number binned into two levels of climatic range – those that were within versus beyond the cold or presumed warm edge, respectively. SE is the standard error. Bold text indicates significance (P < 0.05).

| Model | Model term | Estimate | SE | z-value | P-value |
| --- | --- | --- | --- | --- | --- |
| Optimum (zero inflation) | Intercept | 0.61 | 1.2 | 0.51 | 0.61 |
|  | Temp. range | -42.92 | 30.17 | -1.42 | 0.15 |
|  | **Temp. range<sup>2</sup>** | **31.21** | **13.74** | **2.27** | **0.02** |
|  | Ppt. range | 46.9 | 25.94 | 1.81 | 0.07 |
|  | **Ppt. range<sup>2</sup>** | **-34.68** | **13.03** | **-2.66** | **0.007** |
| Optimum (conditional) | Intercept | -0.83 | 0.53 | -1.55 | 0.12 |
|  | Temp. range | 12.23 | 6.74 | 1.81 | 0.07 |
|  | **Temp. range<sup>2</sup>** | **-25.8** | **5.25** | **-4.92** | **<0.001** |
|  | Ppt. range | 19.49 | 11.33 | 1.72 | 0.08 |
|  | **Ppt. range<sup>2</sup>** | **-44.03** | **8.48** | **-5.2** | **<0.001** |
| Cold edge (zero inflation) | Intercept | -19.31 | 7116.77 | -0.003 | 0.99 |
| Cold edge (conditional) | **Intercept** | **-4.69** | **0.57** | **-8.23** | **<0.001** |
|  | **Cool range (factor)** | **1.37** | **0.35** | **3.93** | **<0.001** |
|  | **Ppt. range** | **-37.54** | **17.46** | **-2.15** | **0.03** |
|  | **Ppt. range<sup>2</sup>** | **-69.89** | **13.11** | **-5.33** | **<0.001** |
| Warm edge (zero inflation) | Intercept | -15.4 | 968 | -0.016 | 0.98 |
| Warm edge (conditional) | **Intercept** | **-2.56** | **0.86** | **-2.97** | **<0.001** |
|  | Warm range (factor) | -0.6 | 0.51 | -1.18 | 0.24 |
|  | **Ppt. range** | **24.94** | **11.26** | **2.22** | **0.02** |
|  | **Ppt. range<sup>2</sup>** | **-24.5** | **7.25** | **-3.38** | **<0.001** |

Species recruited successfully beyond their adult cold edge, further supporting the lagged response hypothesis (Prediction 2; Fig. 5B; Table 1). However, recruitment was reduced beyond the cold edge compared to the cooler half within the range, suggesting that a cold seedling edge is being approached. Nine out of 13 species that had sites beyond their cold edge and potential to recruit in the experiment (i.e., did not exhibit recruitment failure everywhere) successfully recruited beyond their cold edge (recruited beyond cold edge: *Abies grandis, M. aquifolium, M. nervosa, Picea sitchensis, Pinus ponderosa, R. ursinus, T. grandiflora, T. menziesii, and V. parvifolium;* failed to recruit beyond cold edge but recruited within range: *Carex stipata, E. lanatum, Rubus spectabilis, Sambucus caerulea*). However, two species that failed to recruit beyond the cold edge recruited in ≤ 3 plots across the entire experiment (*C. stipata, R. spectabilis*), while all other species with sites beyond their cold edge had higher recruitment success across the experiment (Table S1).

Contrary to the lagged response hypothesis, species successfully recruited within presumed unsuitably warm regions of the adult range (Prediction 3; Fig. 5A; Table 1). Based on the optimal recruitment shift of -0.26, the presumed seedling warm edge was 1 - 0.26 = 0.74 (i.e., shifted 13% from the adult warm edge). Recruitment did not differ between the warmer half of the range (0 - 0.74) and beyond the presumed seedling warm edge (>0.74). All six species with sites beyond the presumed seedling warm edge and the potential to recruit in the experiment (i.e., did not exhibit recruitment failure everywhere) successfully recruited beyond the presumed seedling warm edge (*A. lasiocarpa, A, occidentalis, E. peregrinus, P. engelmannii, Sorbus sitchensis, V. deliciosum*). For a more conservative estimate of recruitment at warm edges in case thermal range shifts are not symmetrical, 3/5 species successfully recruited at and beyond the warmer 5% of their adult range (*A. lasiocarpa, E. peregrinus,* and *V. deliciosum*).

Species successfully recruited beyond their dry and wet edges, through recruitment was limited beyond the wet edge (Fig. S6; Table S2). All seven species with recruitment potential and sites beyond the presumed seedling dry edge (defined as < -0.88 based on the optimal recruitment shift) successfully recruited beyond this dry edge (*M. nervosa, R. ursinus, S. sitchensis, V. parvifolium, T. grandiflora, V. deliciosum, T. grandiflora*). Contrastingly, only two out of eight species with recruitment potential and sites beyond the wet edge successfully recruited beyond the wet edge (recruited beyond wet edge: *A. grandis, P. engelmannii;* failed to recruit beyond wet edge: *A. lasiocarpa, C. stipata, E. lanatum, P. ponderosa, S. caerulea, S. racemosa*).

### Microclimatic buffering hypothesis

Contrary to our predictions for the microclimatic buffering hypothesis, canopy cover had negligible effects on the thermal range position of optimal recruitment or recruitment at either range edge (Prediction 4; optimum: 0.01 ± 0.01, z = -1.00, p = 0.32; cold edge: 0.003 ± 0.02, z = 0.22, p = 0.82; warm edge: 0.04 ± 0.09, z = -0.46, p = 0.64; Table S4). At the microscale, closed canopies in our system had cooler, and less variable temperatures than open canopies, creating the microclimates we assume for the microclimatic buffering hypothesis (Fig. S9 C-F). While open canopies had a cooler thermal seedling optimum compared to closed canopies, consistent with the microclimatic buffering hypothesis, statistical support for this trend was lacking (Fig. 4C; Fig. S10; Table S4).

### Seedling establishment following initial recruitment

Seedling survival results indicate that recruitment patterns identified in previous models are not overturned by countervailing patterns in survival for up to five years (Fig. S11; Table S5). Survival either did not change across thermal ranges (Probability of seedlings surviving to their second year and to the end of the experiment) or showed a similar quadratic relationship as seedling count, peaking in cooler regions of the thermal range (probability of surviving three years). Seedling height increased towards the warm edge, though the effect size was small (0.19 ± 0.07, z = 2.67, p = 0.007; Fig. S11A), suggesting slightly more rapid early growth towards warm range edges.

## Discussion

Findings supported the lagged response hypothesis where apparent range stasis in the North Cascades, USA (Wilson, 2017) masks substantial climatic disequilibrium. Specifically, when seeds were experimentally added (overcoming potential dispersal limitation), we found that many species successfully recruited beyond their leading, cold edge and, on average, the seedling optimum shifted to historically cooler and wetter parts of adult ranges. Meanwhile, recruitment success at warm and dry edges suggests no lagged contractions at the seedling stage at these edges (Fig. 5A). Recruitment was highly limited across our study region, suggesting early establishment constrains species’ ability to track shifting climates, contributing to lagged responses. Contrastingly, we found no evidence supporting the microclimatic buffering hypothesis, but numerical trends were consistent with tree canopy buffering, and the lack of statistical support could reflect low power from low recruitment. In short, our results indicate lagged range expansions at seedling stages towards and beyond cool portions of species ranges, but no lagged contractions at warm, dry edges.

### Lagged climate-driven range shifts

Optimal seedling recruitment shifted from the thermal centre of adults, who recruited in a historically cooler climate, to the cooler (uphill) portion of the adult range as the region has become warmer (Fig. 3; Fig. 4A; Table 1). Adult ranges have recently shifted upslope for many species (Lenoir et al., 2008; Rumpf et al., 2018) and similar mismatches between seedling and adult optima have been observed (Lenoir et al., 2009; but see O’Sullivan et al., 2021) Interestingly, previous work in this system found little evidence of recent shifts in optimal elevation in our study system (Wilson, 2017). We believe this shift may not yet be detected in adults though it is present in seedlings, such that species abundances might be redistributing upslope slowly. Our focal species are long-lived perennials, contributing to the potential for lagged shifts. At the macro-scale of our study system, temperature decreases with elevation, thus indicating upslope distribution shifts as climate warms. Additionally, temperature varies at the microscale across our study system (Chardon et al. *in review*), therefore our findings suggest pockets of changes in abundance within the range as distributions shift upslope.

Focal species recruited beyond their cold edges, indicating ongoing range expansions. However recruitment was reduced beyond the cold edge compared to within the range. This reduced recruitment suggests that, although seedling stages are shifting upward faster than adult stages, dispersal and establishment still lag overall rates of warming. Indeed, recruitment was highly limited across our study region, suggesting early establishment constrains species’ ability to track shifting climates. While we were unable to assess how this reduced seedling recruitment scales up to lifetime fitness, other transplant studies have identified reduced fitness beyond species’ ranges (Hargreaves et al., 2014; Lee-Yaw et al., 2016). We suggest that a lack of observed range expansions might indicate slow ongoing range shifts due to climate lags, creating colonization credits that have not yet been detected at the adult stage at leading, cold edges.

Although seedling responses were consistent with the lagged response hypothesis at the cool region of their ranges, we detected no recruitment failure at warm edges. All six species successfully recruited within the warmer 13% of their adult range and 3/5 recruited up to their adult warm edge, suggesting no climatic mismatches between adult and seedling warm edges. Additionally, we found seedlings exhibited greater growth towards the warm edge (although effects were small), which could increase establishment likelihood towards warm edges. Successful regeneration at the warm edge suggests that these edges are not trailing, as often assumed, and that there is no extinction debt. Warm edges remain suitable for regeneration, despite recent warming possibly due to either a lack of temperature sensitivity or microclimatic buffering by factors other then canopy cover (Chardon et al. *in review*). However, later life stages than what we monitored can pose limiting bottlenecks for population-level replacement at the warm edge in a warmer climate. Additionally, warm, low elevation limits are often considered predominately influenced by species interactions over climate (MacArthur, 1972; Paquette & Hargreaves, 2021), such that non-climatic factors could be more important at warm edges, causing insensitivity to direct effects of temperature increases in warmer portions of the range.

We also found recruitment lags are particularly great in wetter portions of species ranges (Fig 4B; Fig. S5), implying complex responses to warming and drying at seedling stages. Some ranges could be more responsive to precipitation changes than warming (Cahill et al., 2014; Crimmins et al., 2011). Increased recruitment in wetter regions suggests a possible lagged, moisture-driven abundance shift away from drier parts of the range (Fig. S5). However, species recruited at their dry edge, indicating conditions have not yet become too arid for establishment (Fig. S6). Projected precipitation changes in the North Cascades are complex. Historically wetter western slopes are projected for reduced snowpack, more rain, and seasonality changes (Elsner et al., 2010; Mote & Salathé, 2010), while historically drier eastern slopes are projected for an overall precipitation increase, but a temporal shift with drier summers and wetter winters (PCIC, 2021; Stöckle et al., 2009). Thus, inferences about precipitation-driven range shifts depend on which aspects of precipitation are important for establishment. Responses to these fine-scale moisture variations are species-specific in our system (Chardon et al. *in review*). Thus, we are unable to disentangle these microscale moisture drivers in our macroclimatic analysis, but suggest potential distribution shifts from arid to wetter regions within the range. Within our system, precipitation varies along a west-to-east rainshadow gradient and less along an elevation gradient, suggesting potential longitudinal abundance shifts, especially for species that are particularly sensitive to drier conditions.

### Variable species-specific recruitment patterns across the range

Species-specific variation in range shift lags can reflect differences in dispersal, demography, and non-climatic requirements (Alexander et al., 2018). Similar to other studies, we found that climatic mismatches between seedlings and adults are not ubiquitous across species (O’Sullivan et al., 2021; Rabasa et al., 2013) with heterogeneous recruitment patterns among species (Fig. S8). We observed no clear trends based on functional group, although generally low recruitment meant that we lacked the power to explicitly test for trait specific responses across our 25 focal species. However, other studies have found that species traits tend to have low explanatory power in predicting range shifts (Angert et al., 2011; MacLean & Beissinger, 2017). Variable recruitment patterns between species could reflect the differing relative importance of climatic and non-climatic microscale factors for recruitment (Chardon et al. *in review*). For example, *V. deliciosum* had increased recruitment toward and beyond its warm edge. The species might be outcompeted by warm-adapted, low elevation species at later life stages in warmer regions. Similar *Vaccinium* species are competitively inferior in warmer environments (Kudo & Suzuki, 2003). These seemingly idiosyncratic species-level recruitment patterns demonstrate the complexity of climate change responses and can lead to lagged reshuffling of community composition across climatic gradients. Regardless, the community-level pattern elucidates broadscale, ongoing, lagged range shifts and emphasizes the need for multi-species approaches to understand climate change responses.

### No evidence of microclimatic buffering

Microclimatic buffering by canopy cover has the potential to mediate range shifts induced by macroclimatic warming (Greiser et al., 2020; Maclean & Early, 2023). Sheltered, closed canopies can protect species from macroscale warming at warm edges, buffering range contractions. Meanwhile, exposed open canopies can facilitate establishment beyond cold edges (Tourville et al., 2022). Although we did not find evidence that canopy cover affects recruitment across species’ ranges in this way, at least at the early life history stages we monitored (Table S4), our results suggest a possible trend where optimal recruitment shifted further to the cooler portion of the range in more open canopies (Fig. 4C; S6). Thus, there might be a role of canopy cover that we did not have sufficient power to detect as we had limited data in more open canopies. Understanding suitable microsite availability for recruitment of other environmental variables (e.g., soil microbes, moisture, nutrients) across elevation gradients can further elucidate the role of microclimatic buffering in range expansion (Chardon et al. *in review*).

### Study limitations

Our large-scale, multi-species seed addition experiment provided us with many important insights, however, there are limitations to our study and many remaining questions. Because focal species are long-lived, it is unknown whether later demographic transitions enhance or counteract the patterns we found. However, patterns were maintained throughout our five-year experiment (Fig. S11; Table S5). Additionally, there is uncertainty in our calculation of species climatic ranges, particularly for species with limited herbaria records available (Table S1). However, re-running analyses excluding species with < 50 herbaria records (i.e., highest potential for uncertainty in their adult ranges) did not alter our results. Furthermore, since more seeds were added to cooler range positions than warmer, there is a potential bias towards higher recruitment in cooler regions of the range. To test for this, we rarefied the data to an equal number of seed addition plots across thermal ranges. While rarefied datasets were too small for quantitative modelling, we qualitatively assessed rarefied data and confirm optimal recruitment still peaked in cooler regions of the range. Additionally, the optimal recruitment shift between seedlings and adults might, in part, represent local adaptation of species. Recruits followed in this experiment (from locally sourced seeds in northern regions of the temperate forest biome) might be locally adapted to cooler climates and have a lower thermal optimum than the region-wide adult optima we quantified with herbaria records across the temperate forest biome (Ecoregions, 2017). However, we think this is unlikely because we observed recruitment at the warm edge, suggesting locally sourced seeds have similar thermal tolerances to adults. Lastly, many species exhibit annual variation in recruitment (Werner et al., 2020). We monitored for new seedlings for three years to detect delayed recruitment; however, stochastic processes and variability in temporal suitability may, in part, explain why species differ in recruitment success.

## Conclusion

Many species’ ranges appear to remain static throughout recent climate change. We show that experimental seed addition can elucidate upward shifts in performance (in this case seedling recruitment) to cooler, higher elevations for North Cascades montane plant species that are not yet shifting their adult ranges (Wilson, 2017). Either these seedlings will not reach adulthood, or upslope range expansions are occurring at a rate that has yet to be detected amongst adults. Specifically, most species recruited beyond their leading, cold edge and, community-level recruitment optimum shifted to cooler regions. Meanwhile, recruitment at warm and dry edges suggests lower and more arid edges in this system might not be trailing in response to climate warming. Although there is growing evidence that microclimate can buffer range shifts (e.g., Maclean & Early, 2023; Sanczuk et al., 2023), our study suggests range stasis is mostly temporal lags that are disguising species’ vulnerability to climate change. Thus, lagged range shifts imply a future sensitivity of species to climate change that will only become apparent over long time periods or when other factors overcome time-lagged disequilibrium. For example, fire can kickstart range shifts (Wilson, 2017) by facilitating range expansions beyond cold edges and removing relic adult populations at warm edges. Overall, our experiment suggests that adult ranges are not in equilibrium with their current climate, with lagged range shifts likely to occur slowly or in a more punctuated fashion.

## Acknowledgements

We are grateful to C. Lysgaard, L. McBurnie, L. Dorsch, M. Wood, R.K. McCallum, N. Maslowski, A. Cable, O. Rahn, M. Urquhart-Cronish, J. Fowler, J. Rivera-Ordonez, K. Nielson, A. Touch, K. Ertel, K. Gibbs, M. Haenel, J. Hild, R. Klee, E. Lia, A. Kinne, S. Martin-Blangy, T. O’Mara, T. Ota, E. Pletcher, and A. Wall for fieldwork. We thank R. Germain and J. Williams for feedback on earlier versions of this manuscript. A. Kruger and W. van der Bijl provided guidance on bootstrapping R code. This project was funded by the U.S.A. National Science Foundation (to JHRL and AA; grant ID: 1555883), National Science and Engineering Research Council of Canada (Discovery Grant to AA [grant ID: 010335]; CGS-D to KJAG), the University of British Columbia (four-year fellowship and tuition funds to KJAG), and Swiss National Science Foundation (postdoctoral mobility fellowship to NIC [Grant ID: 194331]).

## Data Availability

Upon publication, we will archive our data on Dryad and publish our R code in a public Github repository.

