## supplementary information for "Lagged climate-driven range shifts at species’ leading, but not trailing, range edges revealed by multispecies seed addition experiment"

*Details on study design and study species*

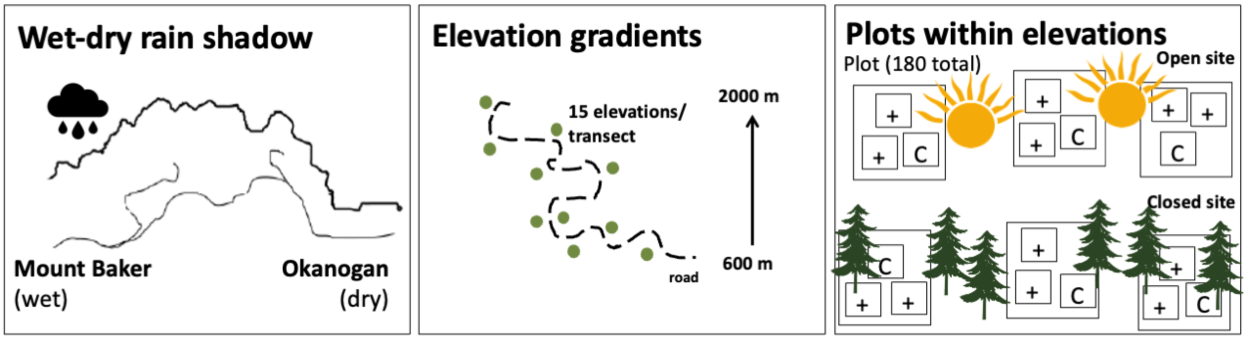

**Figure S1.** Diagram of study design for seed addition experiment. We established one transect each on the wetter west side (Mount Baker National Forest) and drier east side (Rainy Pass; Okanagan National Forest) of a rain shadow gradient on the traditional lands of the Nlaka'pamux, Nooksack, Okanogan, and Methow peoples. 15 elevations per transect were selected. We selected two sites within each elevation: one each with relatively more open and closed canopy cover. Each site had three replicate plots containing three 50 cm x 50 cm quadrats ~20 cm from one another. The three quadrats consisted of a control quadrat (C; no seeds added) and two quadrats with seeds added (+). We split species within the same family into separate quadrats to aid in species identification at the seedling stage. In total, there were 2 transects x 15 elevations x 2 canopy covers x 3 replicate plots x 3 quadrats for 540 quadrats (or 180 plots).

**Table S1.** Description of climatic range and recruitment for each focal species. Lower and upper range edges are, respectively, the 2.5% and 97.5% quantile of mean annual temperature (MAT) or mean annual precipitation (MAP) of herbaria occurrences (http://data.gbif.org/; downloaded on Jan. 31, 2022) from 1981-2010 obtained from Climate NA (Wang et al., 2016).

| Species | Group | Seed source | # herbaria occurrence records | Lower thermal range edge (MAT °C) | Upper thermal range edge (MAT °C) | Lower precipitation range edge (MAP mm) | Upper precipitation range edge (MAP mm) | Seed addition plots recruited (/180) | Sites beyond cold edge | Recruit beyond cold edge | Site in presumed unsuitably warm | Recruit in presumed unsuitably warm |
|---|---|---|---|---|---|---|---|---|---|---|---|---|
| *Abies grandis* | tree | nursery | 14 | 6.13 | 12.14 | 586 | 1908 | 5 | Y | Y | N | |
| *Abies lasiocarpa* | tree | nursery | 107 | -1.77 | 7.08 | 551 | 2425 | 10 | N | | Y | Y |
| *Anemone occidentalis* | forb | local | 28 | 0.37 | 7.16 | 799 | 7416 | 10 | N | | Y | Y |
| *Carex spectabilis* | sedge | local | 114 | -0.7 | 8.47 | 580 | 3076 | 0 | N | | Y | NA |
| *Carex stipata* | sedge | nursery | 104 | 4.86 | 14.3 | 440 | 2022 | 1 | Y | N | N | |
| *Eriophyllum lanatum* | forb | nursery | 442 | 2.5 | 15.2 | 367 | 2822 | 8 | Y | N | N | |
| *Erigeron peregrinus* | forb | local | 54 | -1.17 | 8.37 | 653 | 5405 | 23 | N | | Y | Y |
| *Lupinus latifolius* | forb | both | 161 | 1.9 | 14.5 | 475 | 3588 | 31 | N | | N | |
| *Mahonia aquifolium* | shrub | both | 59 | 4.24 | 13.53 | 367 | 3421 | 8 | Y | Y | N | |
| *Mahonia nervosa* | shrub | local | 75 | 5.89 | 13.18 | 732 | 3593 | 9 | Y | Y | N | |
| *Maianthemum dilatatum* | forb | local | 34 | 3.81 | 11.77 | 717 | 3534 | 0 | Y | NA | N | |
| *Maianthemum racemosum* | forb | local | 272 | 3.13 | 13.3 | 420 | 3381 | 0 | Y | NA | N | |
| *Picea engelmannii* | tree | nursery | 95 | -0.3 | 7.83 | 365 | 1964 | 11 | N | | Y | Y |
| *Picea sitchensis* | tree | nursery | 26 | 5.14 | 11.74 | 579 | 3368 | 7 | Y | Y | N | |
| *Pinus contorta* | tree | nursery | 189 | -0.19 | 11.92 | 441 | 3111 | 0 | N | | N | |
| *Pinus ponderosa* | tree | nursery | 239 | 4.6 | 15.02 | 308 | 2268 | 4 | Y | Y | N | |
| *Rubus spectabilis* | shrub | local | 48 | 3.89 | 12.77 | 526 | 3610 | 3 | Y | N | N | |
| *Rubus ursinus* | shrub | nursery | 103 | 7.51 | 15.17 | 688 | 3301 | 33 | Y | Y | N | |
| *Sambucus caerulea* | shrub | both | 59 | 5 | 13.3 | 311 | 2113 | 2 | Y | N | N | |

| | | | | | | | | | | | | |
|---|---|---|---|---|---|---|---|---|---|---|---|---|
| *Sambucus racemosa* | shrub | both | 215 | 1.4 | 11.39 | 444 | 2623 | 4 | N | | N | |
| *Sorbus sitchensis* | shrub | nursery | 58 | -0.26 | 9.86 | 801 | 4031 | 78 | N | | Y | Y |
| *Tellima grandiflora* | forb | nursery | 45 | 4.97 | 13.6 | 835 | 3803 | 29 | Y | Y | N | |
| *Tolmiea menziesii* | forb | nursery | 45 | 5.87 | 12.68 | 1052 | 3744 | 33 | Y | Y | N | |
| *Vaccinium deliciosum* | shrub | local | 22 | 1.6 | 8.12 | 1180 | 3658 | 24 | N | | Y | Y |
| *Vaccinium parvifolium* | shrub | both | 104 | 5.76 | 12.8 | 782 | 4043 | 43 | Y | Y | N | |

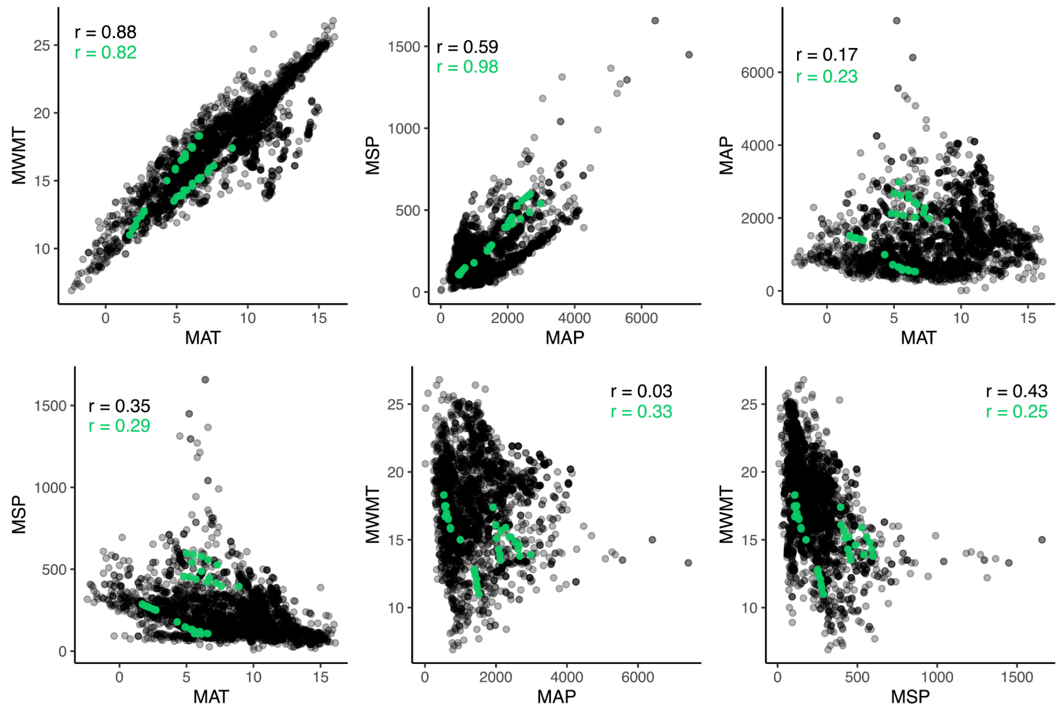

**Figure S2.** Correlations between climatic variables for herbaria records (black) and for seed addition sites (green) amongst mean annual temperature (MAT), mean warmest month temperature (MWMT), mean annual precipitation (MAP), and mean summer precipitation (MSP) from 1981-2010 downloaded from climate NA (Wang et al., 2016). We selected MAT and MAP to respectively represent species thermal and precipitation ranges, which are uncorrelated.

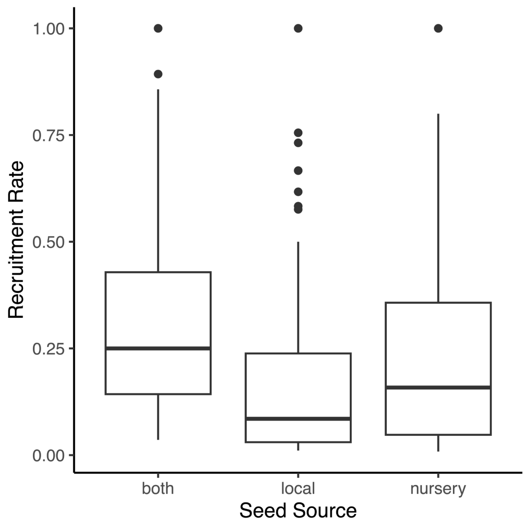

**Figure S3.** Recruitment rate (total recruit counts / number of seeds added) does not significantly vary with seed source (i.e., local source, nursery, or mix of both). We first conducted an ANOVA to test for differences between seed source group mean recruitment rates above 0 with species as a random effect (z-value = 1.008) followed by a Tukey-Kramer posthoc test to identify if group menas varied significantly (local-both P=0.19; both–nursery P = 0.57; local-nursery p = 0.62). Only recruitment rates > 0 are shown.

*Recruitment across precipitation range*

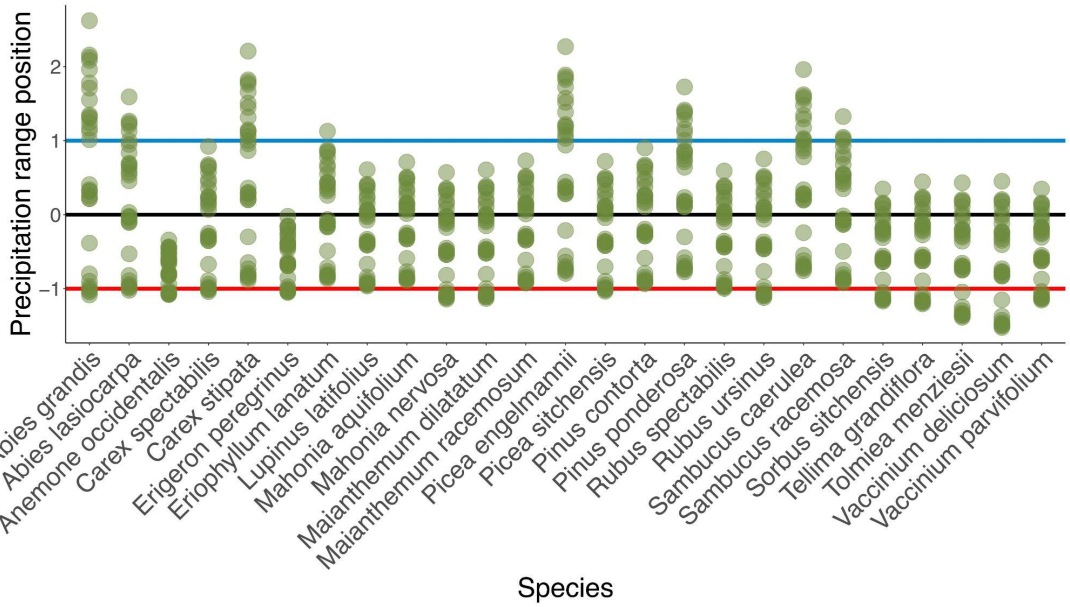

**Figure S4.** Precipitation range position of each study site for each species. Green points indicate a range position of a study site. Blue line is the wet range limit, black line is the precipitation centre, red line is the dry limit. Precipitation range position was quantified using mean annual precipitation data from 1981-2010 (Wang et al., 2016) for herbaria occurrences for each species downloaded from GBIF (http://data.gbif.org/; downloaded on Jan. 31, 2022).

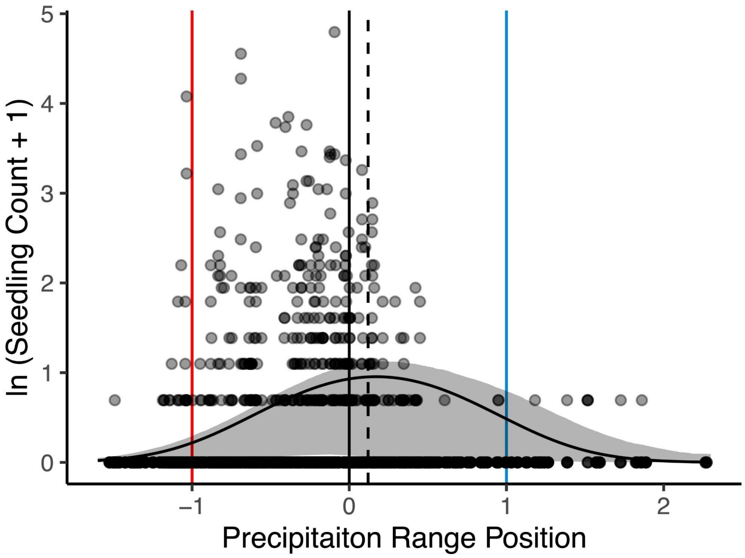

**Figure S5.** Predicted recruitment with precipitation range for the community level with species as a random effect. The black curve is the fitted line from the conditional portion of the zero-inflated model. Shading shows the bias corrected 95% confidence interval around the fitted line derived from a bootstrapping procedure. Points represent the number of recruits for a species in a plot. The cold edge, thermal centre, and warm edge are respectively shown with vertical blue, black, and red lines. The recruitment optimum is shown with a vertical dashed black line.

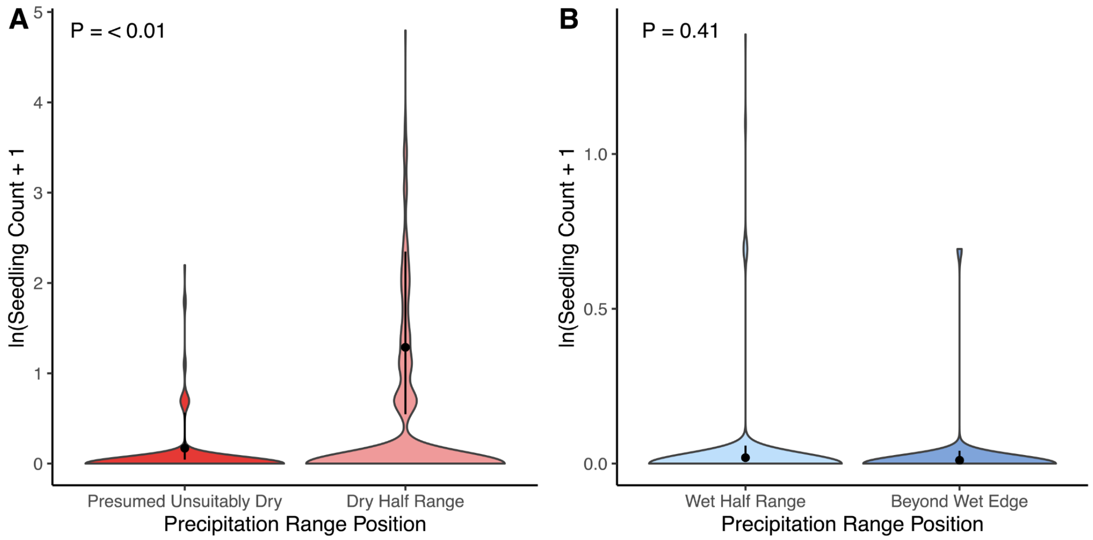

**Figure S6.** Predicted differences in recruitment for (A) presumed unsuitably dry regions of the gradient and the dry half of the range; and (B) the wet half of the range and beyond the wet edge. Species recruited beyond both range edges. The presumed unsuitably dry bin is defined as less than the presumed seedling dry edge (precipitation range position < -0.88, based on the shift in optimal recruitment between adults and seedlings). Dry half of range is a precipitaiton range position -0.88 to 0. Beyond dry edge is a precipitation range position > 1. Wet half of range is a precipitation range position 0 to 1. Shaded violin plot shows the distribution recruits for each species in a given plot. Colour indicates precipitation range bin below the presumed dry edge (dark red), within the drier (light red) or wetter (light blue) half of the range, and beyond the wet edge (dark blue). Black point shows predicted recruitment count, lines show 95% confidence intervals for predicted recruitment.

**Table S2.** Results of zero-inflated generalized mixed linear models testing whether seedling recruitment varies within and beyond species' precipitation range edges as per the lagged response hypothesis. Models compared seedling number binned into two levels of precipitation range – those that were within versus beyond the wet or presumed dry edge, respectively with (temp. range). SE is the standard error. Bold text indicates significance (P < 0.05).

| Model | Model term | Estimate | SE | z-value | P-value |
|---|---|---|---|---|---|
| Dry edge (zero inflation) | Intercept | -12.89 | 158.69 | -0.08 | 0.94 |
| Dry edge (conditional) | **Intercept** | **-3.60** | **0.65** | **-5.52** | **<0.001** |
| | **Dry range (factor)** | **2.65** | **0.51** | **5.19** | **<0.001** |
| | **Temp. range** | **-55.21** | **11.09** | **4.98** | **<0.001** |
| | **Temp. range$^2$** | **-16.27** | **3.85** | **-4.23** | **<0.001** |
| Wet edge (zero inflation) | Intercept | -15.39 | 10283 | -0.001 | 0.99 |
| Wet edge (conditional) | **Intercept** | **-4.53** | **0.77** | **-5.89** | **<0.001** |
| | Wet range (factor) | 0.57 | 0.70 | 0.82 | 0.41 |
| | Temp. range | 8.99 | 10.10 | 0.89 | 0.37 |
| | Temp. range$^2$ | 11.73 | 6.60 | 1.77 | 0.08 |

*Zero-inflated portion of optimal recruitment model*

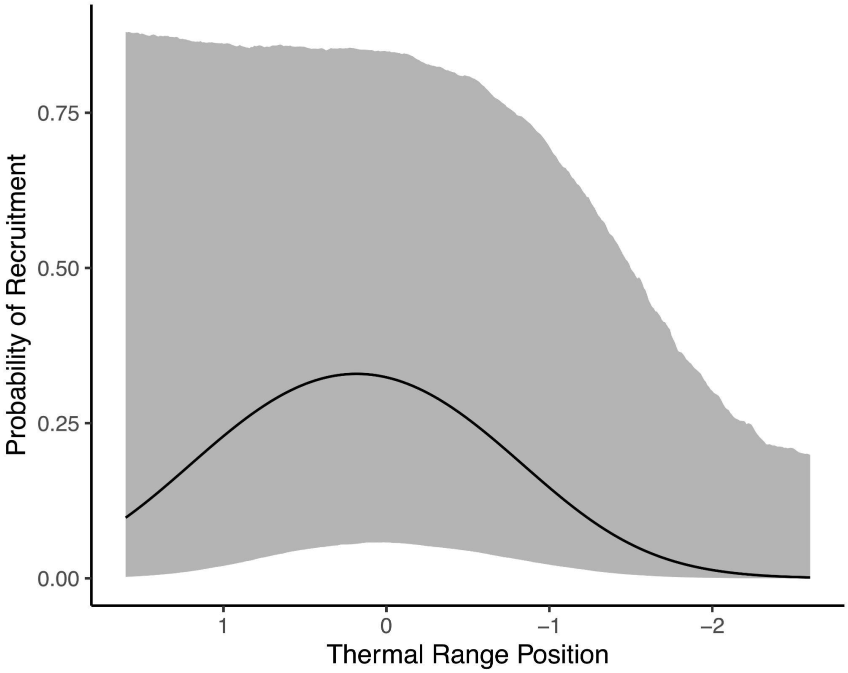

**Figure S7.** The probability of recruitment success did not vary with range position. Prediction for zero-inflation portion of model assessing recruitment success across the thermal range (i.e., probability of any recruitment; see Fig. 3 for conditional model prediction). Solid line is the fitted line from linear models. Gray shading shows the bias corrected 95% confidence intervals around the fitted lines derived from a bootstrapping procedure. Model results summarized in Table 1.

*Species-specific recruitment patterns across the thermal range*

To identify species-specific recruitment patterns across thermal ranges, we ran separate models for the 14 individual species that recruited in at least 8 plots. If species recruited in >23 plots, we constructed negative binomial models with thermal range position and precipitation range position as quadratic fixed effects and site as a random effect. If species recruited in 8-23 plots, we dropped the site random effect so the model could converge. We interpret these reduced species-specific models with caution due to small sample sizes, and simply infer an idiosyncratic response of recruitment across thermal ranges at the species level (Fig. S5; Table S2).

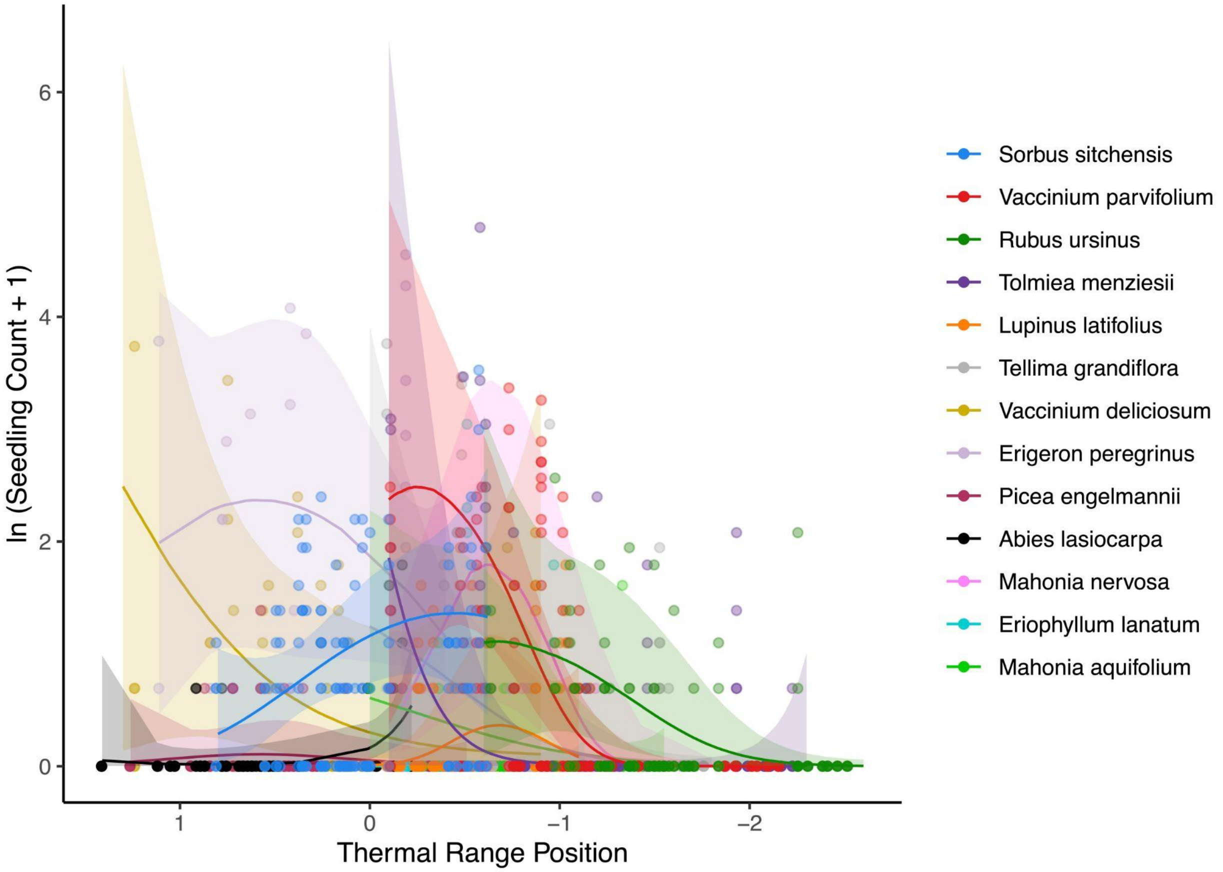

**Figure S8.** Species-specific models results for the 14 species models had wide confidence intervals, limiting our ability to interpret peak recruitment at the species level. The solid lines are the fittings from linear models. Shading shows 95% confidence intervals around fitted lines. Points represent the number of recruits in a plot for each species, denoted by colour.

**Table S3.** Results of species-specific generalized mixed effects linear models testing whether seedling recruitment varies with climatic range. Model complexity varies with sample size (see description above). SE is the standard error. Bold text indicates significance (P < 0.05).

| Species model | Model term | Estimate | SE | z-value | P-value |
|---|---|---|---|---|---|
| *Sorbus sitchensis* (zero inflation) | Intercept | -18.7 | 6004.7 | -0.003 | 0.99 |
| *Sorbus sitchensis* (conditional) | **Intercept** | **-0.68** | **0.26** | **-2.59** | **0.009** |
| | **Temp range** | **-5.84** | **2.91** | **-2.01** | **0.04** |
| | Temp range² | -2.55 | 3.23 | -0.79 | 0.43 |
| | **Ppt. range** | **17.05** | **4** | **4.26** | **<0.001** |
| | **Ppt. range²** | **-10.64** | **3.86** | **-2.76** | **<0.001** |
| *Vaccinium parvifolium* (zero inflation) | **Intercept** | **-1.43** | **0.49** | **-2.89** | **0.004** |
| *Vaccinium parvifolium* (conditional) | **Intercept** | **-4.49** | **1.44** | **-2.12** | **0.002** |
| | **Temp range** | **70.25** | **26.22** | **2.68** | **0.007** |
| | Temp range² | -20.11 | 12.04 | -1.67 | 0.09 |
| | **Ppt. range** | **19.41** | **5.32** | **3.65** | **<0.001** |
| | Ppt. range² | -2.08 | 5.31 | -0.39 | 0.69 |
| *Rubus ursinus* (zero inflation) | Intercept | -19.61 | 9408 | -0.002 | 0.99 |
| *Rubus ursinus* (conditional) | **Intercept** | **-2.59** | **0.57** | **-4.58** | **<0.001** |
| | **Temp range** | **7.24** | **7.24** | **3.13** | **0.002** |
| | Temp range² | 5.52 | 5.52 | -0.92 | 0.36 |
| | Ppt. range | 5.54 | 5.53 | -0.39 | 0.69 |
| | **Ppt. range²** | **8.27** | **8.27** | **-2.37** | **0.02** |
| *Tolmiea menziesii* (zero inflation) | Intercept | -20.89 | 9000 | -0.002 | 0.99 |
| *Tolmiea menziesii* (conditional) | **Intercept** | **-3.94** | **1.14** | **-3.46** | **<0.001** |
| | **Temp range** | **19.87** | **8.41** | **2.36** | **0.02** |
| | Temp range² | 9.66 | 9.28 | 1.04 | 0.3 |
| | Ppt. range | 6.92 | 8.86 | 0.78 | 0.44 |
| | Ppt. range² | -18.02 | 10.63 | -1.7 | 0.09 |
| *Lupinus latifolius* (zero inflation) | Intercept | -19.25 | 10684 | -0.002 | 0.99 |
| *Lupinus latifolius* (conditional) | **Intercept** | **-2.71** | **0.62** | **-4.34** | **<0.001** |
| | Temp range | -9.58 | 6.15 | -1.56 | 0.12 |
| | Temp range² | -10.59 | 7.38 | -1.43 | 0.15 |
| | Ppt. range | 5.27 | 5.26 | 1 | 0.32 |
| | Ppt. range² | -9.13 | 6.36 | -1.44 | 0.15 |
| *Tellima grandiflora* (zero inflation) | Intercept | -12.51 | 7574 | -0.002 | 0.99 |

| | | | | | |
|---|---|---|---|---|---|
| *Tellima grandiflora* (conditional) | **Intercept** | **-1.83** | **0.59** | **-3.11** | **0.002** |
| | **Temp range** | **22.97** | **6.03** | **3.81** | **<0.001** |
| | Temp range² | -2.94 | 6.54 | -0.45 | 0.65 |
| | Ppt. range | -6.57 | 6.44 | -1.02 | 0.31 |
| | **Ppt. range²** | **-30.76** | **9.47** | **-3.25** | **0.001** |
| *Vaccinium deliciosum* (zero inflation) | Intercept | -18.23 | 10415 | -0.002 | 0.99 |
| *Vaccinium deliciosum* (conditional) | **Intercept** | **-2.3** | **0.75** | **-3.1** | **0.002** |
| | **Temp range** | **14** | **7.23** | **1.94** | **0.05** |
| | Temp range² | 3.6 | 6.37 | 0.57 | 0.57 |
| | **Ppt. range** | **17.08** | **6.65** | **2.57** | **0.01** |
| | Ppt. range² | -5.75 | 6.73 | -0.85 | 0.39 |
| *Erigeron peregrinus* (zero inflation) | Intercept | 0.83 | 1.37 | 0.6 | 0.54 |
| *Erigeron peregrinus* (conditional) | Intercept | 1.68 | 1.02 | 1.64 | 0.1 |
| | Temp range | 11.46 | 11.92 | 0.96 | 0.33 |
| | Temp range² | -9.29 | 11.36 | -0.81 | 0.41 |
| | **Ppt. range** | **-13.18** | **6.7** | **-1.9** | **0.05** |
| | Ppt. range² | -23.87 | 14.61 | -1.63 | 0.1 |
| *Picea engelmannii* | **Intercept** | **-2.86** | **0.41** | **-7.07** | **<0.001** |
| | Temp range | 9.49 | 7.72 | 1.23 | 0.22 |
| | Temp range² | -7.46 | 7.29 | -1.02 | 0.31 |
| | Ppt. range | -1.29 | 4 | -0.32 | 0.75 |
| | Ppt. range² | -1.08 | 6.02 | -0.18 | 0.86 |
| *Anemone occidentalis* | Intercept | -2.3 | 2.94 | -0.78 | 0.43 |
| | Temp range | -26.22 | 21.84 | -1.2 | 0.23 |
| | Temp range² | -25.82 | 19.23 | -1.34 | 0.18 |
| | Ppt. range | -19.01 | 13.17 | -1.44 | 0.15 |
| | Ppt. range² | -10.25 | 18.2 | -0.56 | 0.57 |
| *Abies lasiocarpa* | **Intercept** | **-1.91** | **0.81** | **-2.35** | **0.02** |
| | **Temp range** | **-13.93** | **5.95** | **-2.34** | **0.02** |
| | Temp range² | 10.59 | 7.08 | 1.5 | 0.13 |
| | Ppt. range | 0.23 | 7.14 | 0.03 | 0.97 |
| | Ppt. range² | 2.73 | 9.15 | 0.3 | 0.76 |
| *Mahonia nervosa* | **Intercept** | **-7.57** | **2.24** | **-3.38** | **<0.001** |
| | **Temp range** | **89.08** | **37.59** | **2.37** | **0.02** |
| | **Temp range²** | **-36.6** | **15.58** | **-2.35** | **0.02** |

| | | | | | |
|---|---|---|---|---|---|
| | Ppt. range | -27 | 16.21 | -1.67 | 0.1 |
| | **Ppt. range²** | **-41.78** | **17.44** | **-2.4** | **0.02** |
| *Eriophyllum lanatum* | **Intercept** | **-10.29** | **2.97** | **-3.47** | **<0.001** |
| | **Temp range** | **-121.27** | **41.22** | **-2.94** | **0.003** |
| | **Temp range²** | **39.2** | **13.83** | **-2.83** | **0.005** |
| | **Ppt. range** | **-53.31** | **17.92** | **-2.97** | **0.003** |
| | **Ppt. range²** | **61.2** | **23.35** | **2.62** | **0.009** |
| *Mahonia aquifolium* | **Intercept** | **-3.78** | **1.05** | **-3.6** | **<0.001** |
| | Temp range | 17.56 | 13.44 | 1.31 | 0.19 |
| | Temp range² | -3.41 | 9.83 | -0.35 | 0.73 |
| | Ppt. range | -29.53 | 20.31 | -1.45 | 0.15 |
| | Ppt. range² | -30.02 | 24.23 | 1.24 | 0.22 |

*Microclimatic buffering hypothesis results*

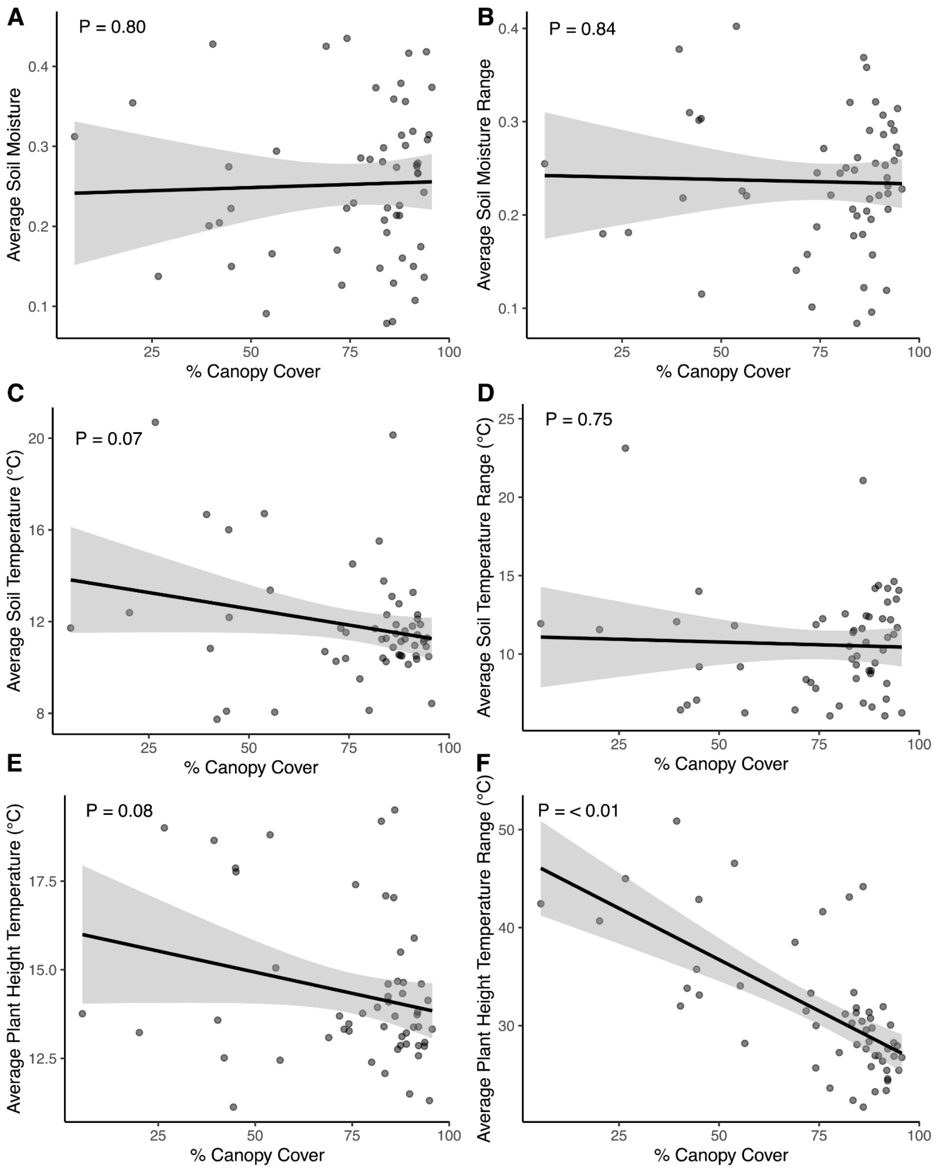

**Figure S9.** Microclimatic changes with canopy cover in our study system. Temperature is cooler and less variable in increasingly closed canopies, as assumed for the microclimatic buffering hypothesis. Soil moisture does not change with canopy cover. We recorded soil moisture and soil and plant height temperature (15 cm aboveground) throughout the growing season at each site from May - September 2022 (using TMS-4 data loggers, TOMST, Prague Czech Republic; Wild et al., 2019). We calculated growing season variables from these data loggers to estimate growing season mean daily average and mean daily range (daily maximum - daily minimum) each for soil moisture (A,B), soil temperature (C,D) and plant height temperature (E,F). See Chardon et al. *in review* for additional details on microclimatic data collection. Each point represents a study site. Lines and shading show prediction and 95% confidence intervals from simple linear regression including the microclimatic response variable and canopy cover.

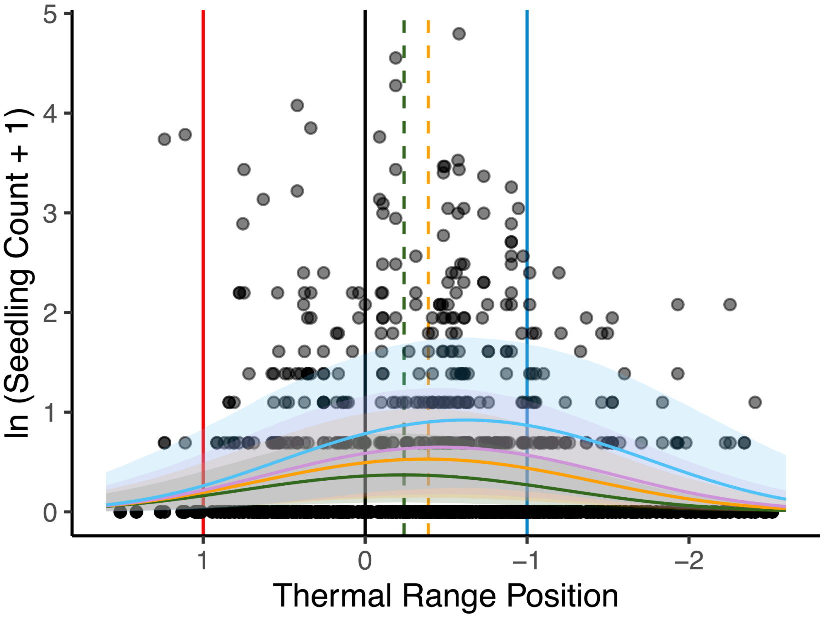

**Figure S10.** Predicted relationship between recruitment and thermal range position for different canopy covers. There was a non-significant interaction term for recruitment between canopy cover and thermal range position (P = 0.32), but relatively more open canopies had optimal recruitment in the cooler part of adult thermal ranges than closed canopies. 88% canopy cover (mean canopy cover of closed canopy sites; green), 64% canopy cover (mean of open sites; orange), 50% canopy cover (purple), and 25% canopy cover (blue) fitted lines are shown (solid curves). Shading shows the bias corrected 95% confidence interval around the fitted lines derived from a bootstrapping procedure. Points represent the number of recruits for a given species in a plot. The optima for 88% and 64% canopy cover (i.e., canopy cover optimum histogram visualized in Fig. 4C) is shown in green and orange dashed vertical lines. Note that we have less data for relatively more open canopies, thus 50% and 75% canopy cover optima are not visualized in the Fig. 4C histogram. The cold edge, thermal centre, and warm edge are respectively shown with vertical blue, black, and red lines. Model results are summarized in Table S3.

**Table S4.** Results of zero-inflated, generalized, mixed effects linear models testing whether seedling recruitment varies with canopy cover and climatic range as per the microclimatic buffering hypothesis for 25 species in the North Cascades, WA. All models include species and plot nested within site as random effects (results not shown). SE is the standard error. Bold text indicates significance ($P < 0.05$).

| Model | Model term | Estimate | SE | z-value | P-value |
|---|---|---|---|---|---|
| Cold edge (zero inflation) | Intercept | -12.58 | 245.72 | -0.05 | 0.96 |
| Cold edge (conditional) | **Intercept** | **-4.22** | **0.65** | **-6.45** | **<0.001** |
| | **Temp. range (factor)** | **1.23** | **0.48** | **2.53** | **0.01** |
| | **Ppt. range** | **-41.16** | **17.65** | **-2.33** | **0.02** |
| | **Ppt. range$^2$** | **-68.73** | **13.11** | **-5.24** | **<0.001** |
| | Canopy | -0.02 | 0.01 | -1.32 | 0.19 |
| | Canopy:temp. range | 0.003 | 0.02 | 0.22 | 0.82 |
| Warm edge (zero inflation) | Intercept | -18.44 | 4378.99 | -0.01 | 0.99 |
| Warm edge (conditional) | Intercept | -1.99 | 1.29 | -1.54 | 0.12 |
| | Temp. range (factor) | -1.03 | 1.12 | -0.93 | 0.36 |
| | **Ppt. range** | **24.64** | **11.45** | **2.15** | **0.03** |
| | **Ppt. range$^2$** | **-24.01** | **7.34** | **-3.27** | **0.001** |
| | Canopy | -0.05 | 0.09 | -0.55 | 0.58 |
| | Canopy:temp. range | 0.04 | 0.09 | -0.46 | 0.64 |
| Optima shift (zero inflation) | Intercept | -0.32 | 1.15 | -0.28 | 0.78 |
| | **Temp. range** | **-33.27** | **17.11** | **-1.94** | **0.05** |
| | Temp. range$^2$ | 19.76 | 13.17 | 1.50 | 0.13 |
| | **Canopy** | **0.04** | **0.02** | **1.96** | **0.05** |
| Optima shift (conditional) | Intercept | -0.88 | 0.62 | -1.43 | 0.15 |
| | **Temp. range** | **22.05** | **9.19** | **2.40** | **0.02** |
| | **Temp. range$^2$** | **-28.22** | **5.91** | **-4.78** | **<0.001** |
| | Region | -0.66 | 0.35 | -1.87 | 0.06 |
| | Canopy | 0.02 | 0.01 | 1.60 | 0.11 |
| | Canopy:temp. range | 0.01 | 0.01 | -1.00 | 0.32 |

*Seedling establishment following initial recruitment*

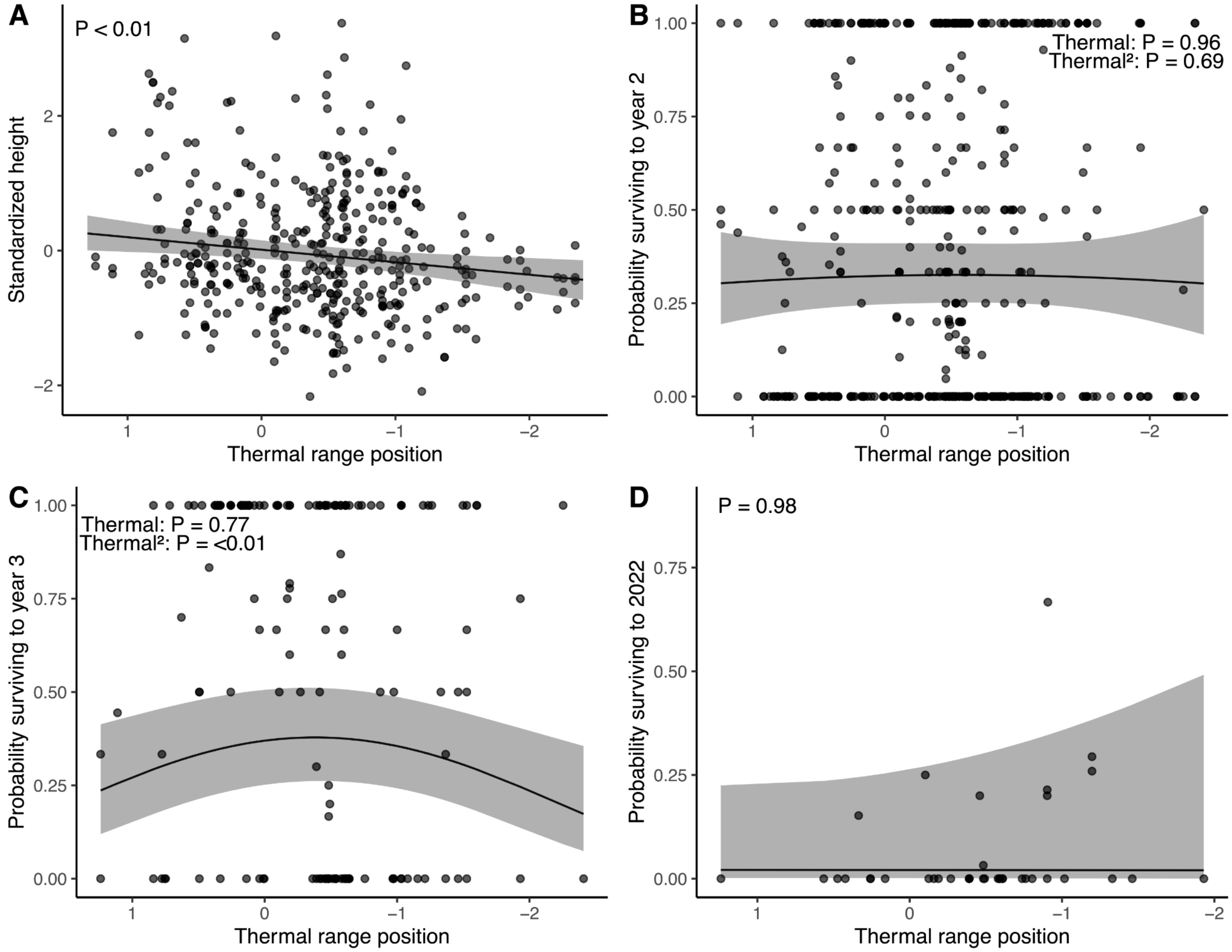

**Figure S11.** Predicted relationship across the thermal range for seedling (A) height, (B) survival to the second year, (C) survival to the third year, and (D) survival to the end of the experiment (up to five years). Solid line shows predicted fit from linear models. Gray shading shows 95% confidence intervals. Each point is either the mean standardized height or proportion of seedlings surviving for a given species in a plot. Height increases towards warmer thermal range positions. Probability of seedling survival either did not change with thermal range position (B, D), or peaked in cooler regions of the range (C), similar to the seedling count model. Model results summarized in Table S4.

**Table S5.** Results of generalized mixed effects linear models testing whether seedling survival and height varied with climatic range for 25 species in the North Cascades, WA. All models include plot nested within site as random effects (results not shown). SE is the standard error. Bold text indicates significance ($P < 0.05$).

| Model | Model term | Estimate | SE | z-value | P-value |
|---|---|---|---|---|---|
| Survive to 2nd year | **Intercept** | **-0.76** | **0.16** | **-4.65** | **<0.001** |
| | Temp. range | -0.06 | 1.51 | -0.04 | 0.97 |
| | Temp. range$^2$ | -0.48 | 1.23 | -0.39 | 0.69 |
| | **Ppt. range** | **-10.28** | **1.23** | **-3.56** | **<0.001** |
| | **Ppt. range$^2$** | **-7.44** | **3.30** | **-2.26** | **0.02** |
| | **Year germinated** | **0.50** | **0.12** | **4.22** | **<0.001** |
| Survive to 3rd year | Intercept | -0.42 | 0.25 | -1.66 | 0.10 |
| | Temp. range | 0.44 | 1.53 | 0.29 | 0.77 |
| | **Temp. range$^2$** | **-3.03** | **1.07** | **-2.84** | **0.004** |
| | Ppt. range | -2.94 | 2.77 | -1.06 | 0.29 |
| | Ppt. range$^2$ | 4.20 | 2.39 | 1.75 | 0.08 |
| Survive to 2022 | **Intercept** | **-3.85** | **1.44** | **-2.67** | **0.008** |
| | Temp. range | 0.01 | 0.46 | 0.03 | 0.98 |
| | Ppt. range | 4.61 | 3.42 | 1.35 | 0.18 |
| Height 1st year | Intercept | 0.01 | 0.08 | 0.18 | 0.85 |
| | **Temp. range** | **0.19** | **0.07** | **2.67** | **0.007** |
| | Ppt. range | -0.02 | 0.11 | -0.2 | 0.84 |

*References*

Wang, T., Hamann, A., Spittlehouse, D., & Carroll, C. (2016). Locally Downscaled and Spatially Customizable Climate Data for Historical and Future Periods for North America. *PLOS ONE*, *11*(6), e0156720. https://doi.org/10.1371/journal.pone.0156720

Wild, J., Kopecký, M., Macek, M., Šanda, M., Jankovec, J., & Haase, T. (2019). Climate at ecologically relevant scales: A new temperature and soil moisture logger for long-term microclimate measurement. *Agricultural and Forest Meteorology*, *268*, 40–47. https://doi.org/10.1016/j.agrformet.2018.12.018
